# Temporal specificity and heterogeneity of the fly immune cells’ transcriptional landscape

**DOI:** 10.1101/2019.12.20.871301

**Authors:** Pierre B. Cattenoz, Rosy Sakr, Alexia Pavlidaki, Claude Delaporte, Andrea Riba, Nacho Molina, Nivedita Hariharan, Tina Mukherjee, Angela Giangrande

## Abstract

Immune cells provide defense against the non-self, however recent data suggest roles well beyond innate immunity, in processes as diverse as development, metabolism and tumor progression. Nevertheless, the heterogeneity of these cells remains an open question. Using bulk RNA sequencing we find that the *Drosophila* immune cells (hemocytes) display distinct features in the embryo, a closed and rapidly developing system, compared to the larva, which is exposed to environmental and metabolic challenges. Through single cell RNA sequencing we identify fourteen hemocyte clusters present in unchallenged larvae and associated with distinct cellular processes e.g. proliferation, phagocytosis, metabolic homeostasis and humoral response. Finally, we characterize the changes occurring in the hemocyte clusters upon wasp infestation that triggers the differentiation of a novel cell type, the lamellocyte. This first molecular atlas provides precious insights and paves the way to study the biology of the *Drosophila* immune cells in physiological and pathological conditions.

## Introduction

The innate immune response has made the object of intense investigation in *Drosophila melanogaster*, as this model shows mechanisms that are conserved throughout evolution, from pattern recognition molecules to immune molecular cascades (Akira et al., 2006; Kleino and Silverman, 2014). Given the importance of innate immunity in a variety of physiological and pathological processes including tumour progression (Ratheesh et al., 2015), the current challenge is to characterize immune cell heterogeneity and identify specific hemocyte populations. This is the aim of the present work.

Three classes of hemocytes have so far been identified: plasmatocytes (PL), crystal cells (CC) and lamellocytes (LM) (Honti et al., 2014). PL are the most abundant cell type and are responsible for the main functions of the hemocytes: phagocytosis, secretion of extracellular matrix proteins (ECM), signalling molecules and antimicrobial peptides (AMPs) (Baer et al., 2010; Basset et al., 2000; Ferrandon et al., 2004; Gold and Bruckner, 2015; Sears et al., 2003; Yasothornsrikul et al., 1997). The CC account for less than 5% of the total hemocyte population with distinctive crystals inside them that are composed of prophenoloxidases (PPO) (Rizki and Rizki, 1959). These enzymes are released in large quantity upon wounding and are key component for the melanization process (Rizki and Rizki, 1959). The LM are flat and large cells that only appear upon challenge. They are considered activated immune cells (Gold and Bruckner, 2015) that arise through PL trans-differentiation or from a mitotic dedicated precursor (Anderl et al., 2016).

In the embryo, the hemocytes contribute to the clearance of apoptotic cells and the deposition of ECM-related molecules including Peroxidasin (Pxn) and Viking (Vkg) (Nelson et al., 1994; Yasothornsrikul et al., 1997). By the larval stage, the organism interacts with the external environment and responds to metabolic and oxidative stress as well as to infection or injury related stimuli. The hemocytes must therefore adapt to these new, highly demanding, settings. In addition, while during embryogenesis the hemocytes are highly motile and patrol the whole organism, during the larval life a large fraction of them, called resident hemocytes, colonize segmentally repeated epidermal-muscular pockets in which cell proliferation is enhanced (Makhijani et al., 2011). Upon wounding, septic infection or infestation by parasitic wasps, the resident hemocytes are mobilized and enter in circulation to reach the site of the immune challenge (Dragojlovic-Munther and Martinez-Agosto, 2012; Owusu-Ansah and Banerjee, 2009). Thus, hemocyte localization adapts to homeostatic and challenged conditions.

We here characterize the transcriptional changes occurring during development and the different types of hemocytes present in the larva. Comparing the bulk RNA sequencing data allows us to define stage-specific features: hemocytes contribute to the shaping of the tissues and are glycolytic whereas larval hemocytes show a strong phagocytic potential and a metabolic switch toward internalization of glucose and lipid and beta-oxidation. The single cell RNA sequencing (sc RNA seq) assays allow us to identify fourteen clusters of larval PL and to assign specific molecular and cellular features, including nutrient storage, proliferative potential, antimicrobial peptide production and phagocytosis.

Finally, as a first characterization of the immune response at the single cell level, we assess the transcriptional changes induced by infestation by the parasitic wasp *Leptopilina boulardi*, one of the most studied cellular immune pathways. The wasp lays eggs in the *Drosophila* larva and triggers hemocyte proliferation as well as LM differentiation (Markus et al., 2009), with subsequent encapsulation of the wasp egg and its death through the increase of the levels of reactive oxygen species (ROS). The sc RNA seq sequencing assay identifies two LM populations, a mature one with a strong glycolytic signature, and a population that expresses both LM and PL features, likely originating through trans-differentiation (Anderl et al., 2016).

The response to wasp infestation involves the embryonic hemocytes that differentiate from the procephalic mesoderm (1^st^ wave of hematopoiesis) (Tepass et al., 1994), as well as the hemocytes that originate from the lymph gland, the site of the 2^nd^ hematopoietic wave. While in not infested (NI) conditions, the lymph gland histolyses and releases hemocytes in circulation during the pupal life, upon wasp infestation (WI), it undergoes precocious histolysis so that both lymph gland and embryo derived hemocytes populate the larva (Banerjee et al., 2019; Bazzi et al., 2018; Letourneau et al., 2016). Our single cell RNA sequencing assay identifies the same number of PL clusters as that observed in normal conditions, showing that the PL from the first and from the second hematopoietic wave share the same features.

In sum, this work characterizes the transcriptional changes occurring during hemocyte development and the hemocyte populations present in the *Drosophila* larva. It also provides the molecular signature and the initial characterization of the larval hemocyte repertoire as well numerous novel markers in NI and in WI conditions. These first bulk and single cell RNA seq data pave the way to understand the role of the immune system in development and physiology.

## Result

### Comparing the bulk transcriptomes from embryonic (E16) and larval (WL) hemocytes

In the embryo, insulated from most immune challenges by the eggshell, the hemocytes main functions are developmental. They clear the organism from apoptotic bodies issued from organogenesis and secrete extracellular components. In the larva, the hemocytes display new properties to respond to the microorganism-rich environment in which they grow. To identify the changes occurring in the hemocytes during development, we compared the hemocytes’ transcriptomes from mature, stage 16 embryos (E16) and from wandering larvae (WL).

The comparison shows 3396 genes significantly up-regulated in E16 and 1593 up-regulated in WL (**Figure 1A**, data in **Supplemental Table S1**). Most PL markers such as Hemese (He) (Kurucz et al., 2003), Singed (Sn) (Zanet et al., 2009), Eater (Kocks et al., 2005), Hemolectin (Hml) (Goto et al., 2001), Serpent (Srp) (Shlyakhover et al., 2018), Nimrod C1 (NimC1, also called P1) (Kurucz et al., 2007), Croquemort (Crq) (Franc et al., 1996) and Pxn (Nelson et al., 1994) are strongly expressed at both stages but enriched in WL hemocytes (**Figure 1C**). CC markers are also present in the transcriptome: Pebbled (Peb) and Lozenge (Lz) are detected at relatively low levels, in agreement with the small number of CC in the E16 and WL hemolymph (Rizki and Rizki, 1959). The CC specific markers PPO1 and PPO2, on the other hand, are among the genes expressed at the highest levels, highlighting their key function and the sharp specialization of the CC (Binggeli et al., 2014).

**Figure 1:**
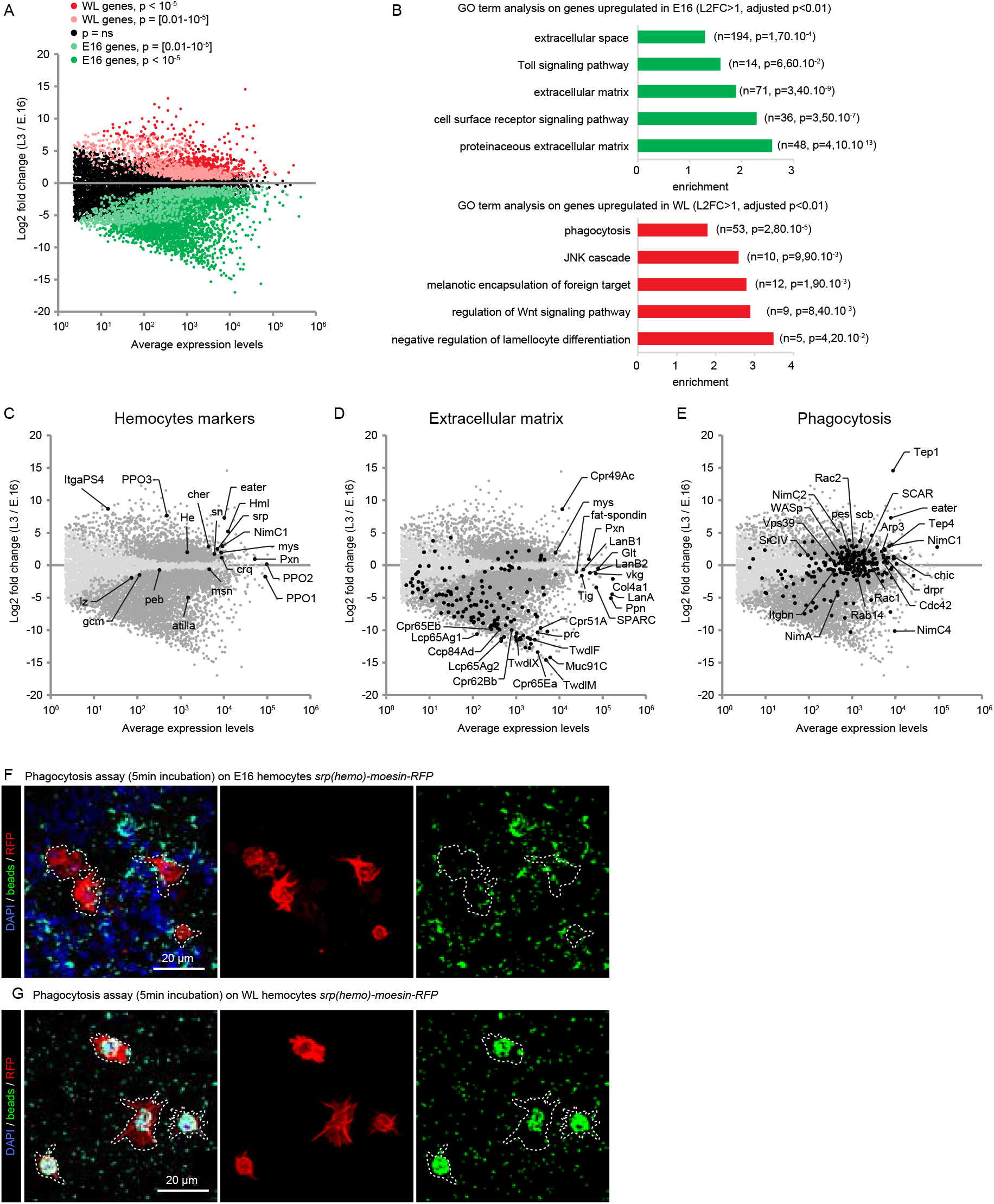
Comparison of the transcriptomes from embryonic stage 16 and wandering L3 hemocytes. **A)** Scatter plot comparing the transcriptome of hemocytes from stage 16 embryos (E16) and wandering 3^rd^ instar larvae (WL). The average levels of gene expression are plotted on the x-axis and the Log2 Fold Change WL/E16 on the y-axis. The p-value of the fold change is indicated with the following color code: not significant in black, enriched in the embryo, with a p-value <10^-5^ in green and with a p-value [0.01-10^-5^] in light green, enriched in the larva, with a p-value <10^-5^ in red and with a p-value [0.01-10^-5^] in light red. The data comparing these two datasets are available in **Supplemental Table S1.** **B)** Gene Ontology (GO) term enrichment analysis in the E16 (in green) and WL (in red) hemocytes. The histograms show the fold enrichment for a subset of significant GO terms (full list in **Supplemental Table S1**), the number of genes associated with the GO term and the p-value of the GO term enrichment are indicated in brackets. **C-E)** Scatter plots highlighting in black subsets of known genes expressed in hemocytes (**C**) or genes associated with the GO terms extracellular matrix **(D)** and phagocytosis **(E)**. Light gray dots indicate genes that are not expressed in a significantly different manner between E16 and WL. Dark gray dots indicate genes that are expressed in a significantly different manner. **F-G)** Confocal images of phagocytosis assay on E16 **(F)** and WL hemocytes **(G)** *srp(hemo)-moesin-RFP*. The beads (in green) are phagocytosed by the hemocytes (in red). The WL hemocytes show greater phagocytic capacity compared to the embryonic ones after 5 min of exposure. Full stacks are displayed and the scale bars represent 20 μm. Related to **Supplemental Figure S1, S2, Table S1, S4.**

Surprisingly, most LM markers such as Myospheroid (Mys or L4) (Irving et al., 2005), Misshapen (Msn) (Braun et al., 1997), Cher (or L5) (Rus et al., 2006) and Atilla (or L1) (Honti et al., 2009) were also detected at significant levels in the hemocytes from both stages, suggesting that they are expressed at basal levels in normal hemocytes and are strongly induced in LM. Finally, Gcm is involved in hemocyte development in the early embryo (stage 8-10) (Bernardoni et al., 1997) and is no longer expressed by E16 (Bazzi et al., 2018). Accordingly, Gcm transcripts are barely detected in E16 and WL transcriptomes (levels < 40 normalized read count). Overall, these data prove the efficiency of the experimental design to purify hemocytes.

### Embryonic hemocytes express ECM components

We next carried out a GO term enrichment analysis on the genes up-regulated in either population (|Log2 fold change WL/E16| > 1, adjusted p-value < 0.01) (**Supplemental Table S1**). The E16 hemocytes display a striking enrichment for genes coding for extracellular matrix components (ECM) (**Figure 1B,D**). Out of 162 genes coding for ECM proteins, 138 are enriched in E16 hemocytes. To confirm the expression pattern of the ECM genes, we compared these data with two *in situ* hybridization databases (Berkeley Drosophila Genome Project (Hammonds et al., 2013; Tomancak et al., 2002; Tomancak et al., 2007) and Fly-FISH (Lecuyer et al., 2007; Wilk et al., 2016)) (**Figure S1D,E**). Most genes for which we could find data are specifically expressed in hemocytes in the embryo (**Figure S1D,E**).

The expression/secretion of specific ECM compounds by the hemocytes during embryonic development was previously described. The integrins alphaPS1 (Mew) and Mys and the integrin ligand Tiggrin (Tig) are secreted by the hemocytes at the level of muscle insertion to stabilize strong attachment between the cells (Bunch et al., 1998; Fogerty et al., 1994). The laminins LanA, LanB1, LanB2 and Wb are secreted by the hemocytes for them to migrate efficiently throughout the embryo (Sanchez-Sanchez et al., 2017). Pxn and the collagens Vkg and Col4a1 secretion by the hemocytes is essential for the condensation of the ventral nerve cord (Olofsson and Page, 2005). Finally, SPARC is produced by the hemocytes and is necessary for basal lamina assembly (Martinek et al., 2008). All these compounds are expressed at similar range in embryo and in larva (**Figure 1D**), suggesting that the role of the hemocytes in secreting these ECM compounds is preserved throughout development. In addition, we identified 21 other genes associated with the ECM term and highly up-regulated in the embryo (Log2FC < −3, pvalue < 0.01 in **Table S1**). This suggests additional, embryo-specific pathways for the deposition of the ECM.

Finally, E16 hemocytes are specifically enriched for a large group of ECM compounds described as constituent of the cuticle: 23 Tweedles (Twdl), 56 Cuticular Proteins (Cpr and Ccp), 9 Larval Cuticle Proteins (Lcp), 9 Mucins (Muc) (**Figure 1D**). Previous reports indicate that embryonic cuticle deposition by the epithelial cells is induced by ecdysone (Charles, 2010; Chavez et al., 2000; Maroy et al., 1988). The transcriptome data indicate that the Ecdysone receptor is expressed in E16 hemocytes. Future investigations will reveal the role of hemocytes in cuticle deposition and whether the expression of the cuticle genes in hemocytes is induced by ecdysone signaling.

### Larval hemocytes express specific scavenger receptors

The GO terms enriched in WL compared to E16 hemocytes highlight phagocytosis and, to a lower extent, signaling pathways involved in the immune response (JNK and Wnt) (**Figure 1B,E, Supplemental Table S1**).

Among the genes involved in phagocytosis, is a large panel of genes coding for transmembrane phagocytic receptors involved in pathogen recognition, such as the Nimrod family (Eater (Kocks et al., 2005), NimC1 (Kurucz et al., 2007) and NimC2), several scavenger receptors (Sr-CIV, He (Kurucz et al., 2003), Draper (Drpr) (Cuttell et al., 2008; Hashimoto et al., 2009), Peste (Philips et al., 2005)) as well as the integrins Scab (alpha-PS3) and Integrin beta-nu (Itgbn) (Nonaka et al., 2013). Noteworthy, the E16 embryonic hemocytes are specifically enriched for NimC4 (also called Simu), a receptor of the Nimrod family that is involved in the phagocytosis of apoptotic bodies (**Figure 1E**) (Kurant et al., 2008; Roddie et al., 2019).

The WL hemocytes are also enriched for Opsonins. These secreted molecules bind to the pathogens and promote their phagocytosis by the macrophages. Tep1 and Tep4 (Dostalova et al., 2017; Haller et al., 2018) are among the genes expressed at the highest levels in WL hemocytes and Tep1 presents the strongest enrichment. Most of the genes involved in phagosome formation are also enriched at this stage: Arp3, Rac1, Rac2, SCAR, WASP, Chic and Cdc42. Finally, genes involved in phagosome maturation (Rab14) and phagolysosome formation (Vps39) are enriched as well (**Figure 1E**).

The scavenger receptors and the opsonins cover a large panel of pathogens (For review see (Melcarne et al., 2019)), indicating an overall switch for hemocytes function from apoptotic body scavenging and cuticle production at embryonic stages to pathogen scavenging at the WL stage. Since it was previously shown that the hemocytes present in the embryo are able of phagocytosis (Tan et al., 2014; Vlisidou et al., 2009), we compared the phagocytic capacity of hemocytes from E16 and WL and exposed them to fluorescent beads. The results clearly show that the larval hemocytes phagocytose faster and more than the embryonic ones (**Figure 1F-G**).

Thus, the transcriptome analysis reveals a clear shift in the function of the hemocytes during development, from building the ECM and the cuticle to adopting a defense profile against immune challenge.

### Metabolic shift between embryonic and larval hemocytes

The transcriptome data also reveal that the larval hemocytes are most likely internalising and metabolising lipids through the beta oxidation pathway to generate acetyl CoA and drive the TCA cycle **(Figure S2A-C)**. This notion is supported by the up-regulation of genes encoding lipid scavenging receptors and the down-regulation of genes involved in lipid biosynthetic (TAG) pathway **(Figure S2A,B)**.

The down-regulation of genes involved in glycolysis, mainly phosphofructokinase and Pyruvate dehydrogenase **(Figure S2B)**, implies that the larval hemocytes do not rely on this process to drive the TCA-cycle. The transcriptional down-regulation of gluconeogenic genes (phosphoenolpyruvate carboxykinase and fructose 1, 6 bisphosphatase) suggests the absence of gluconeogenesis in these cells. However, a significant up-regulation of the Glut1 sugar transporter suggests active uptake of glucose by the larval hemocytes. The down-regulation of glycolytic genes downstream of G6P and up-regulation of genes of the pentose phosphate pathway (PPP) implies that the internalized glucose could be potentially used to generate pentose sugars for ribonucleotide synthesis and redox homeostasis through the generation of NADPH. Corroborating this observation is also the strong up-regulation of redox homeostatic enzymes **(Figure S2A)**.

In contrast to the larval hemocytes, the E16 hemocytes are glycolytic and do not rely on oxidative metabolism **(Figure S2B)**. This is supported by the strong up-regulation of a key glycolytic enzyme, lactate dehydrogenase essential for the conversion of pyruvate to lactate. Furthermore, these cells do not metabolise lipids, as enzymes of the beta oxidation pathway are transcriptionally down-regulated.

### Generation of single cell RNA seq datasets from NI and WI larvae

The *Drosophila* larva contains PL and CC that are resident or in circulation. Upon wasp infestation, the LM are produced from precursors (Anderl et al., 2016) or by PL transdifferentiation (Stofanko et al., 2010). To obtain a comprehensive repertoire of the hemocyte populations present in the larva, we generated two single cell libraries on the hemocytes from not-infested WL (NI dataset) and from WL infested by the parasitoid wasp *L. boulardi* (WI dataset).

The resident and the circulating hemocytes were collected from pools of 20 female larvae. The libraries were produced using the Chromium single cell 3’ mRNA seq protocol (10x Genomics). The NI library contains 7606 cells (mean read per cell = 37288; median genes per cell = 959) and the WI library 8058 cells (mean read per cell = 32365; median genes per cell = 1250). The libraries were merged to cluster the hemocytes presenting similar expression profiles using the Seurat toolkit (Butler et al., 2018; Stuart et al., 2018) (**Figure S3A**). Subclustering was then applied to refine the grouping of the cells leading to the identification of 16 clusters of hemocytes (**Figure S3A’, C-D’’**). The identity of each cluster was assigned using the list of known markers for the CC (Lz, Peb, PPO1, PPO2), for the LM (Mys, Msn, Cher, Atilla, ItgaPS4, PPO3) and for the PL (Sn, Pxn, Hml, Eater, NimC1, Crq, He, Srp) (**Figure S3B**). Of note, the single cell data confirm that the LM markers Mys, Msn, Cher and Atilla detected in the bulk RNA seq on WL are expressed in PL and enriched in LM (**Figure S3B**).

We identified 13 clusters of PL and 1 cluster of CC in both the NI and the WI larvae and 2 clusters of LM specifically found in the WI larvae. The name of each cluster corresponds to the name of one of the main markers or biological features (**Figure 2A,B**). Importantly, all cells analyzed in these datasets present known hemocyte markers, which indicates a high purity of the samples.

**Figure 2:**
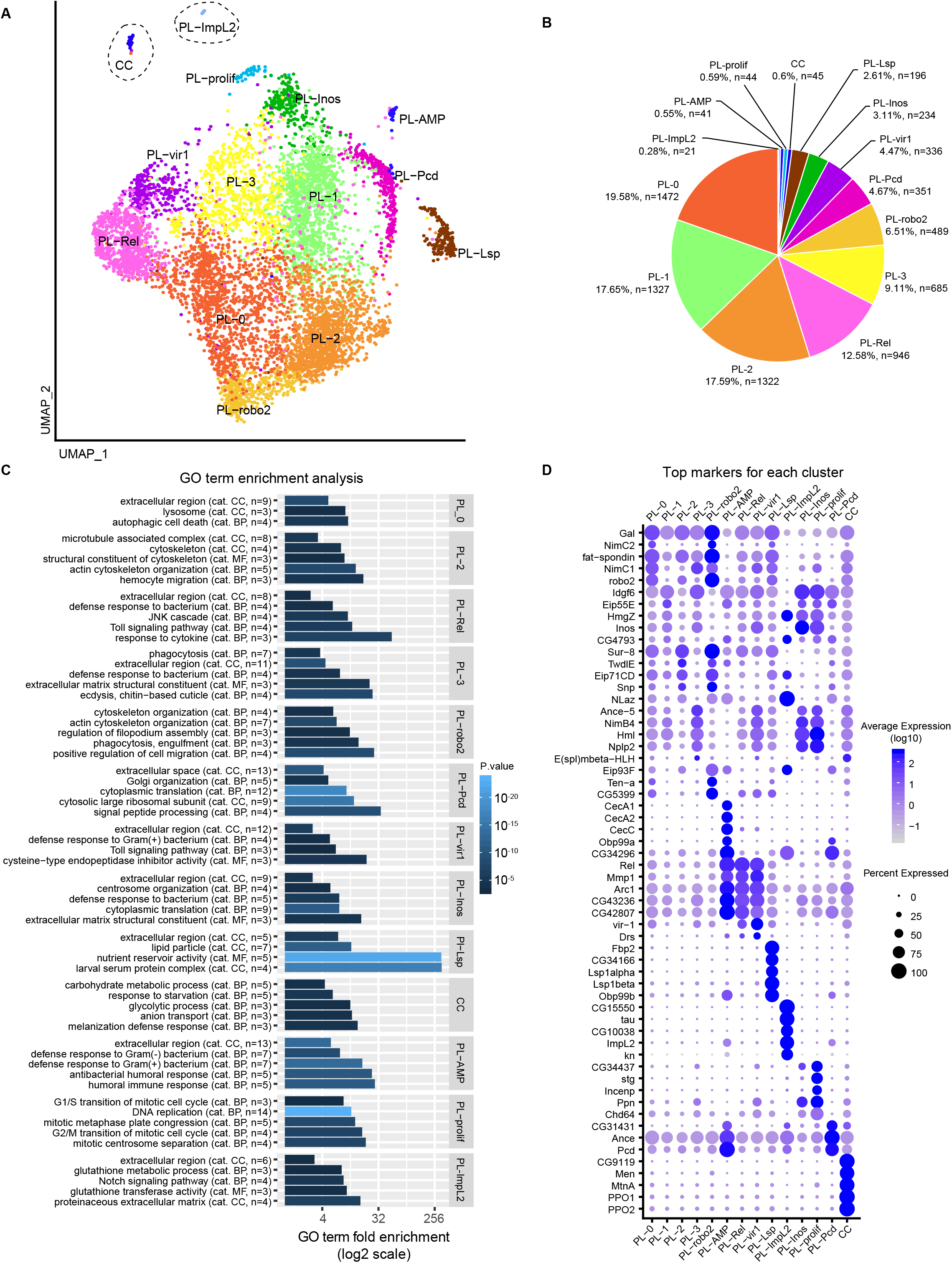
Identification of the hemocyte populations from WL by single cell RNA seq (10x genomics). **A)** UMAP projection representing the 14 clusters of cells identified in the hemocyte pools from *OregonR* WL (NI dataset). **B)** Number of cells and proportion of each cluster in the NI dataset. **C)** Bar graph displaying the GO terms enriched in specific clusters (subset from **Supplemental Table S3**). The x-axis shows the enrichment of the GO term, the color gradient (gradient from black to light blue) indicates the p-value, the number of genes and the GO term category (CC: cellular compound, BP: biological process, MF: molecular function) are indicated between brackets. **D)** Dot plot representing the expression levels (gradient of purple levels) and the percentage of cells (size of the dots) that express the top 5 markers of each cluster. Related to **Supplemental Figure S3, S4, Table S2 and S3**.

### Characterization of the transcriptomic profile in normal conditions

Following the identification of the clusters, our first aim was to characterize the properties of the clusters in the NI dataset. Thus, we carried out GO term analyses on the genes enriched in each of them (**Table S2, Figure 2C**) using DAVID (Huang da et al., 2009). In addition, to estimate whether the clusters are enriched/specifically localized in the circulating or in the resident compartment, we performed qPCR assays on hemocytes from each compartment and targeting the strong markers of the different clusters (**Figure 3A**).

**Figure 3:**
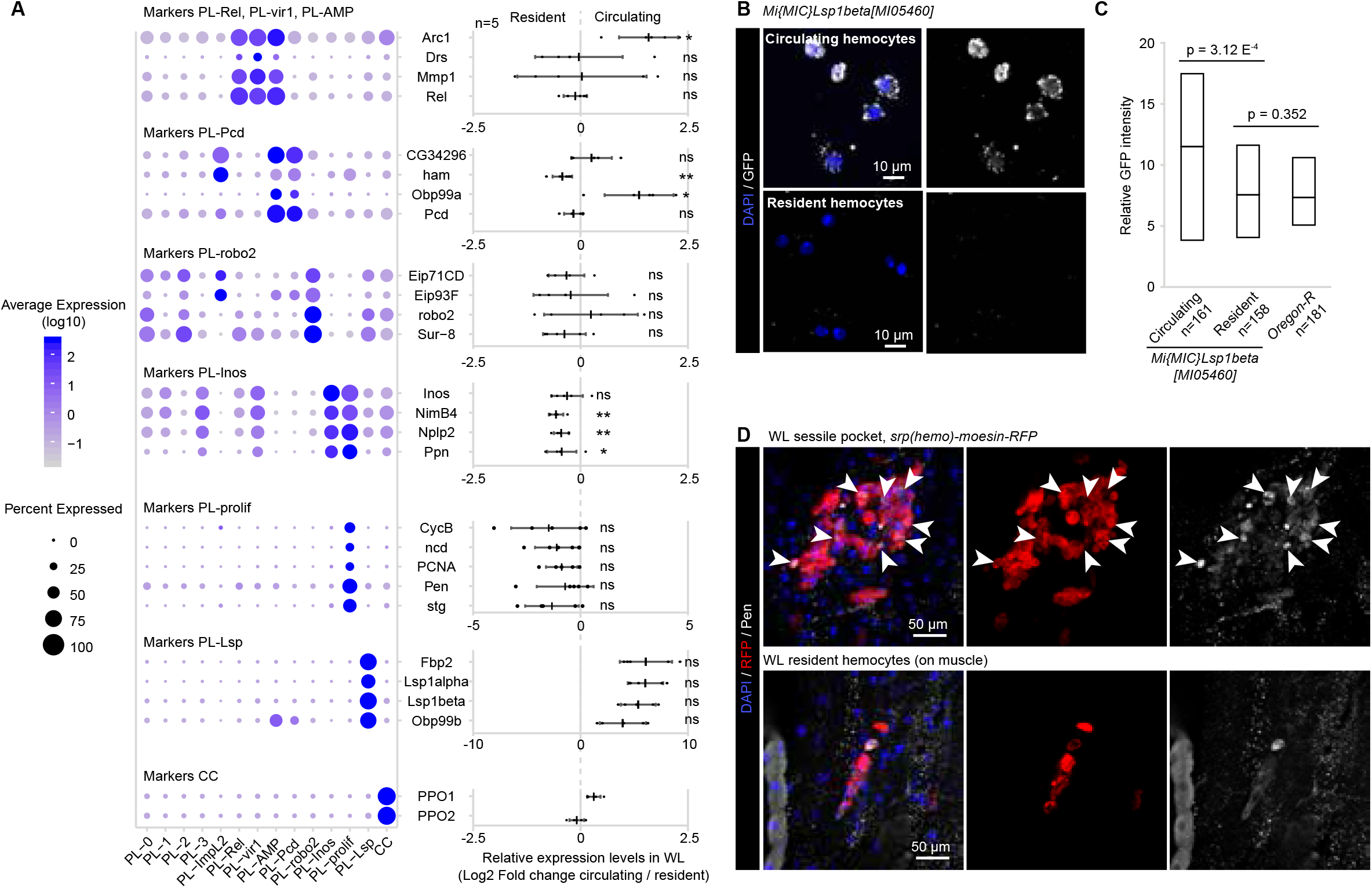
Localization of the NI hemocytes clusters. **A)** Identification of the position (circulating/resident) of the clusters within the larva by qPCR. The dot plot on the left panel indicates the distribution of each marker across all clusters (as in **Figure 2D**), the graph on the right panel, indicates the Log2 of the ratio between the expression level in the circulating versus the resident compartment. Positive values indicate an enrichment in the circulating compartment and negative values indicate an enrichment in the resident compartment. The experiment was carried out on 5 independent replicates (pool of 10 larvae per replicate). The p-values are estimated by bilateral paired student test and indicated as follow: ns = non-significant (>0.05), “*” = p[0.05 – 0.01[, “**” = p[0.01-0.001[, “***” = p<0.001. **B)** Confocal images of circulating and resident hemocytes (top and bottom panels, respectively) from *Mimic-Lsp1beta-MI05460* WL, which express Lsp1beta tagged with GFP. The immunolabeling was done with anti-GFP (in gray) and the nuclei were marked with DAPI (blue). Full stacks are displayed, the left panels show the overlay of DAPI and GFP, the right panels show the GFP alone. The scale bars represent 10 μm. **C)** Quantification of the GFP intensity in circulating and resident hemocytes from *MiMIC-Lsp1beta-MI05460* larvae. The values obtained on the *OregonR* hemocytes indicate the level of the background. The number of cells included in the analysis is displayed on the x-axis label and were quantified from two independent preparation, p-values were estimated after variance analysis with bilateral student test for equal variance. **D)** Confocal images of resident hemocytes located around the oenocytes (Makhijani et al., 2011) or along the muscles (top and bottom panels, respectively) from WL *srp(hemo)-moesin-RFP*, that express moesin tagged with RFP in hemocytes. The immunolabeling was done with anti-RFP (in red), anti-Pendulin (Pen, in gray) and the nuclei were marked with DAPI (blue). Full stacks are displayed, the left panels show the overlay of DAPI, Pen and RFP, the middle panels show RFP alone and the right panels show Pen alone. Arrow heads in the top panels indicate cells that co-express RFP and Pen. The scale bars represent 50 μm. Related to **Supplemental Figure S5, Table S6**.

The PL clusters PL-0, PL-1, PL-2 and PL-3, encompass more than 60% of all the hemocytes. These clusters express most of the PL markers (**Figure S4A**) but do not display strong distinct signatures (avg_logFC of the strongest markers < 0.93, **Figure 2D**). The 9 remaining PL clusters present very specific molecular signatures in addition to the main PL markers (**Figure 2D**) and are organized at the periphery of the 4 clusters on the graphical representation generated with the UMAP dimension reduction technique (Becht et al., 2018) (**Figure 2A**).

### PL-Rel

The PL-Rel cluster includes 12.6% of the total hemocyte population, with more than 100 strong markers (**Table S2**), most of which are involved in the immune responses (**Figure 2C**). The cluster expresses the main transcription factors of the Imd pathway (*i.e*. Relish, Rel) and of the Toll pathway (*i.e*. Dorsal, Dl), Cactus (Cact) as well as the secreted protein PGRP-SA (Govind, 1999; Valanne et al., 2011; Zhai et al., 2018) **(Table S2)**. In addition, it expresses proteins associated with the JNK pathway such as Jra and Puc (Martin-Blanco et al., 1998; Zheng et al., 2017). The qPCR on resident and circulating hemocytes indicates that PL-Rel hemocytes are present in both compartments (**Figure 3A**).

### PL-vir1

The PL-vir1 cluster contains 4.5% of the total hemocytes and expresses the same strong markers than PL-Rel, including Rel, Jra and Puc (**Figure 2D, Table S2**). Compared to PL-Rel, it displays an up-regulation of the marker of viral infection Vir1 (Dostert et al., 2005), the predicted peptidase Ance-5 and the peptide Nplp2, that is secreted upon immune challenge (Verleyen et al., 2006). PL-vir1 presents also same GO terms than PL-Rel, associated with the defense response to bacterium and Toll signaling pathway (**Figure 2C**). The qPCR suggests that PL-vir1 is present in both resident and circulating compartment.

### PL-robo2

The PL-robo2 cluster represents 6.5% of the total hemocyte population. It does not express unique markers (**Figure 2D**), but displays a strong enrichment for GO terms related to migration and phagocytosis (**Figure 2C**). It expresses the actin-regulatory protein Enable (Stedden et al., 2019; Tucker et al., 2011) and transmembrane proteins that participate to the migration of multiple cell types. It also presents the strongest up-regulation of the phagocytic receptors Crq and Drpr (Franc et al., 1999; Manaka et al., 2004). Of note, Crq is the main receptor involved in lipid scavenging and is a major actor of the induction of the inflammatory response to high fat diet initiated by the hemocytes (Woodcock et al., 2015). In addition, PL-robo2 is enriched in the lipid droplet associated protein Jabba involved in the regulation of lipid metabolism (McMillan et al., 2018). Together, these data suggest that the PL-robo2 hemocytes serves as sensor of lipid levels in the hemolymph. PL-robo2 is present in both circulating and resident compartments (**Figure 3A**).

### PL-Pcd

The PL-Pcd cluster contains 4.7% of the total hemocyte population and is mostly linked to translation and Golgi organization (**Figure 2C**). Half of the proteins present in the GO term enriched functions are ribosomal proteins (3 RpS including Rps3 and 11 RpL), while the others are related with Golgi (Gmap, Ire-1, Sec23) (Friggi-Grelin et al., 2006; Norum et al., 2010; Zacharogianni et al., 2011), which suggests either that translation is higher in this cluster compared to other clusters or that the ribosomal proteins, high in this cluster, act on the immune response in an uncanonical fashion. Notably, studies in mammals indicate that ribosomal protein like RPS3 selectively modulates the target genes of NF-κB, the orthologue of Rel (Zhou et al., 2015). This cluster also expresses specifically the pterin Pcd involved in amino acid metabolism, the peptidase Ance involved in proteolysis and two uncharacterized genes CG31431 and CG34296 (**Figure 2D**).

The qPCR data highlight one marker (Ham) enriched in the resident compartment and one marker (Obp99a) enriched in circulation. These ambiguous results may be due to the fact that the markers are expressed in other clusters as well (PL-AMP and PL-ImpL2).

### PL-AMP

The PL-AMP cluster (0.5% of the hemocytes) presents strong similarities with PL-Rel, as they share the same markers related to the Imd pathway (**Figure 2D**), but distinguishes itself by the expression of the antimicrobial peptides (AMP) Cecropin A1 (CecA1), Cecropin A2 (CecA2), Cecropin C (CecC), Attacin-A (AttA), Attacin-B (AttB) and Attacin-D (AttD) **(Figure 2D**, **Table S2**) as well as of the markers of PL-Pcd cluster Pcd, Ance, CG31431 and CG34296 (**Figure 2D**). The AMP are usually induced and secreted primarily by the fat body after septic wounds that triggers the Imd pathway (Govind, 1999; Zhai et al., 2018). These data suggest that PL-AMP represents activated hemocytes. The expression of CecC, CecA1 and CecA2 were not detected by qPCR in the resident and circulating hemocytes. This is likely due to the low representation of these cells combined with the small size of the Cecropin transcripts (less than 400nt) that prevents the optimal design of primers and affects the PCR efficiency. Therefore, we could not conclude on the localization of the PL-AMP hemocyte in the larvae.

### PL-Inos, PL-prolif

The clusters PL-Inos and PL-prolif express several common markers and represent approximately 3% and 0.6% of the total hemocyte population, respectively (**Figure 2B**).

The PL-Inos cluster is enriched in GO terms associated with multiple functions including the response to bacteria, the ECM, cytoplasmic translation and centrosome organization.

The PL-prolif cluster is specifically characterized by genes involved in mitosis (**Figure 2C**, **Table S2**). For example, genes coding for Klp61F, Klp67A and Ncd linked to mitotic centrosome separation (Gandhi et al., 2004; Sharp et al., 1999) as well as Ncd80 and Nuf2 that are part of the NCD80 complex are all up-regulated. Like their mammalian counterparts, these proteins are essential for the mitotic metaphase plate congregation (Przewloka et al., 2007). Finally, PL-prolif hemocytes express two cyclins (CycB and CycE), transporters (Pen) and cell cycle related enzymes (String (Stg), the Cyclin dependent kinases Cdk1 and Cdk2), which are linked to the G2/M and G1/S transition, respectively (Kussel and Frasch, 1995; Yuan et al., 2016).

The qPCR assay indicates that the clusters PL-Inos and PL-prolif are enriched in the resident compartment (**Figure 3A,D**). This is concordant with a previous analysis indicating that the resident hemocytes are more proliferative than the circulating ones (Makhijani et al., 2011).

### PL-Lsp

The PL-Lsp cluster contains approximately 3% of the total hemocyte population and is strongly associated with the GO terms larval serum protein complex, nutrient reservoir activity and lipid particle (**Figure 2C**, **Table S2**). This is due to the expression of the larval serum proteins (LSP), which serve as nutrient pool that will be used during metamorphosis (Telfer and Kunkel, 1991). The PL-Lsp cluster also expresses the receptor responsible for the incorporation of the LSPs (i.e. Fbp1), as well as proteins associated with lipid transport (Rfabg) (Burmester et al., 1999; Kutty et al., 1996; Massey et al., 1997) and the odorant binding protein Obp99b that is considered as a storage protein (Handke et al., 2013). These proteins are usually described as secreted by the fat body, suggesting shared features and role in metabolism between this tissue and PL-Lsp.

The qPCR on resident and circulating PL reveals that the PL-Lsp hemocytes are mostly in circulation (**Figure 3A)**. This localization is supported by labeling of two independent LSP transgenic reporters (**Figures 3B,C**, **S5A-C**).

### PL-Impl2

The PL-ImpL2 cluster comprises less than 0.3% of the hemocytes, but displays the most distinctive molecular signature (49 genes presenting an enrichment avg_logFC > 1, **Table S2**). This includes genes involved in glutathione metabolism (GstD1, GstD3, GstE12) and specific transcription factors (Ham, Kn, Antp, Eip93F, Noc and ElB) (**Figure 2C, S5D,E**).

Many of these markers are usually associated with the post signaling center (PSC) that regulates the differentiation of LM within the lymph gland (Benmimoun et al., 2015; Crozatier et al., 2004; Mandal et al., 2007). These data suggest that the PL-ImpL2 cluster defines PSC-like cells present outside of the lymph gland in a still unknown location. The low abundance of these cells renders them hard to track in the larva, however the GstD reporter *GstD-LacZ* (Sykiotis and Bohmann, 2008) labels subsets of WL hemocytes that may correspond to the PL-ImpL2 cluster (**Figure S5E**).

### CC

The CC cluster expresses the two well-known CC markers PPO1 and PPO2 (Dudzic et al., 2015). In addition, we identify new potential markers: Metallothionein A (MtnA), Malic enzyme (Men) and CG9119. Further analysis of the GO terms highlights the response to starvation (Lipin (Lpin), Mthl10) as well as enzymes essential for the biosynthesis of proteoglycans (Sugarless, sgl) and glucose homeostasis (Pfk, 6-phosphofructo-2-kinase (pfrx), Aldolase (ald)) (Dudzic et al., 2015; Enzo et al., 2015; Flowers et al., 2007; Hacker et al., 1997; Sung et al., 2017; Ugrankar et al., 2011; Wong et al., 2019). This suggests that the CC do not operate with the same metabolic pathway as the majority of the WL hemocytes, for which the bulk RNA seq data indicate a lipid based metabolism (**Figure S2**).

Finally, GO terms related to ‘extracellular region’ are enriched recurrently in almost every cluster (**Figure 2C**). Further analyses of all genes expressed in the NI dataset and associated with the term extracellular region show that each cluster expresses different proteins (**Figure S4B**).

### Characterization of the molecular pathways active in the clusters

To identify which regulatory networks characterize the clusters, we performed a regulon analysis using SCENIC (Aibar et al., 2017) (**Figure 4A**), which determines the regulon(s) active in single cells in three steps. First, covariation of the expression levels of the genes is estimated in the single cell dataset. Then, each group of genes displaying covariation is screened for common cis-regulatory motifs targeted by transcription factors present in the group. This defines groups of genes regulated by a specific transcription factor (= regulon). Finally, the activity of each regulon is estimated in each cell of the dataset (Aibar et al., 2017). Of note, this analysis was done independently of our initial clustering based on the expression levels. The clustering of the WT NI cells based on the regulons is highly comparable to the expression based clustering, with an overlap between the two clustering approaches, estimated with the Rand Index (Rand, 1971) of 0.83 (**see the method section**). This overlap highlights the robustness of our initial clustering.

**Figure 4:**
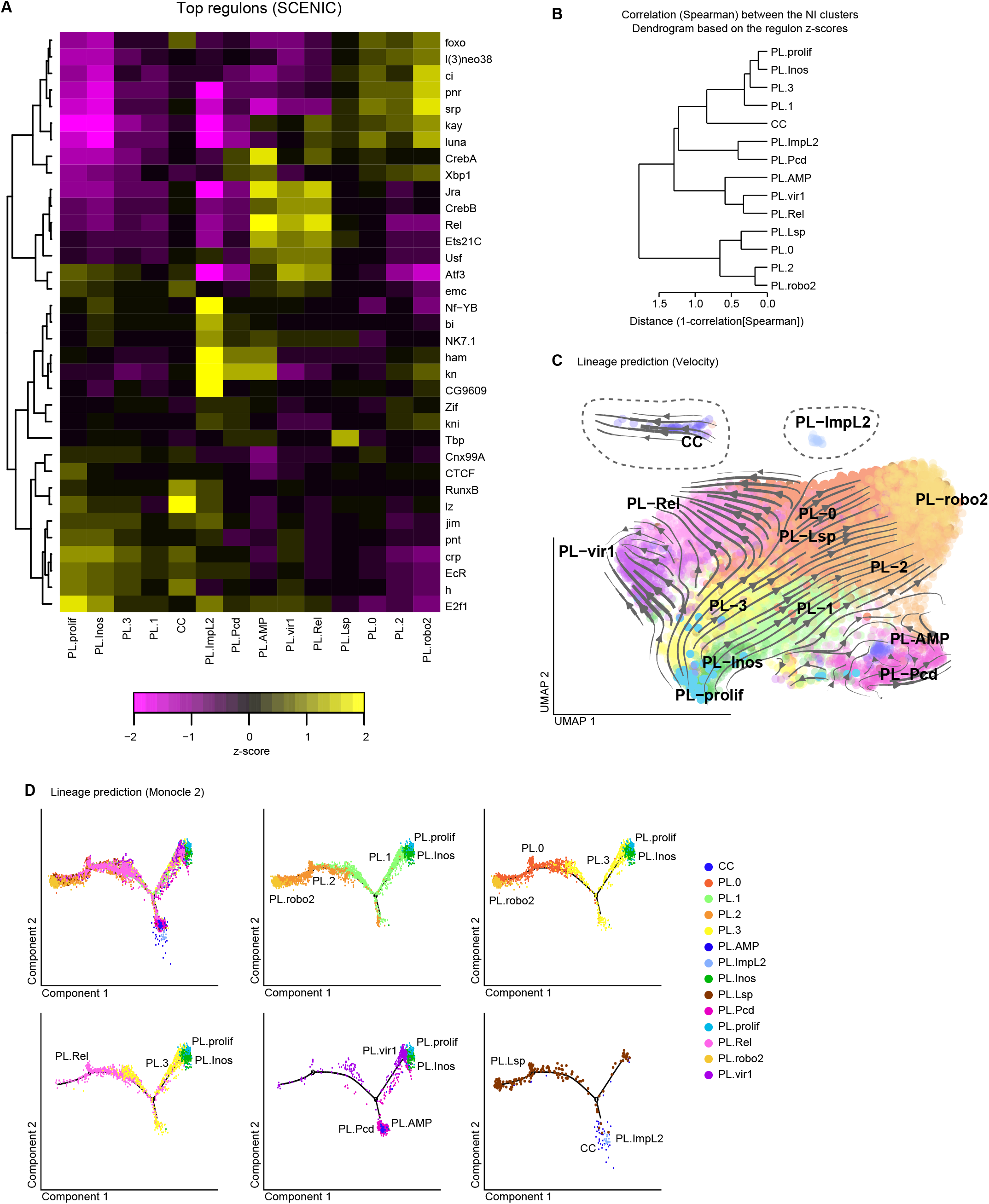
Identification of the cluster specific molecular pathways and of the filiation between the NI clusters. **A)** Heatmap representing the enrichment (z-score) for the top 5 regulons of each cluster determined with SCENIC. The dendrogram on the left side of the panel indicates the correlation between the regulons across the dataset. The enrichment per cluster is indicated with a gradient from magenta (z-score < 0, repression of the regulon in the cluster) to yellow (z-score > 0, activation of the regulon in the cluster). **B)** Dendrogram representing the distance among the clusters. The tree was built on the correlation (Spearman) calculated on the regulon matrix from the NI dataset. **C)** Lineage prediction using RNA Velocity. The arrows and lines on the UMAP predict the “direction” taken by the cells of each cluster, based on the comparison between the levels of mRNA and pre-mRNA. Note that the clusters CC and PL-ImpL2 (dashed lines) have been moved from their original position to fit in the graph. **D)** Single cell trajectory reconstructed with monocle 2 on the NI dataset. The first panel shows the overlap of all clusters and subsequent panels show restricted number of clusters for which the RNA velocity analysis suggested a filiation.

The SCENIC analysis allowed us to associate specific regulons to each cluster (**Figure 4A**). As a proof of concept, the analysis highlighted a positive correlation between the regulon lz and the CC. Lz is a Runx transcription factor essential for the differentiation of these cells (Lebestky et al., 2000).

-The PL-Rel, PL-AMP and PL-vir1 clusters are characterized by the regulons Jra (JNK cascade), Rel (IMD pathway) and Ets21C (which cooperates with the JNK pathway). The regulon Activating Transcription Factor 3 (ATF3) involved in anti-viral response in mammals (Labzin et al., 2015) is specific to PL-vir1 and PL-Rel, whereas the regulon CrebA regulating the secretory pathway is specific to the PL-AMP cluster, which expresses most of the antimicrobial peptides.

-The PL-prolif cluster and the closely related cluster PL-Inos (**Figure 4B**) are enriched for the regulons EcR and E2F1 that are involved in the regulation of hemocyte proliferation (Sinenko et al., 2010).

-PL-Lsp is associated with the Tbp regulon that is involved in the canonical transcriptional machinery. Such enrichment may indicate a higher rate of transcription, which would be concordant with the function of this cluster in producing storage proteins in preparation for pupariation.

-PL-robo2 is enriched for the regulons of the GATA factors Pnr and Srp, which is known to regulate the expression of scavenger receptors in the hemocytes (Shlyakhover et al., 2018; Valanne et al., 2018).

-At last, PL-ImpL2 is enriched in Ham, Kn, CG9609 and Nf-YB. Ham was shown to limit amplifying division in neural stem cells (Eroglu et al., 2014). CG9609 is a zinc finger transcription factor poorly described, expressed mostly in ovaries (Robinson et al., 2013) and Nf-YB regulates cell death and proliferation (Ly et al., 2013).

### Developmental links between the different hemocyte populations in NI animals

The GO term and the regulon analyses reveal distinct functions and properties for specific hemocyte clusters. The identification of the proliferative cluster prompted us to ask whether there is a filiation among the clusters and, if so, to define their hierarchical organization. We adopted two distinct strategies to predict the filiation: RNA Velocity (La Manno et al., 2018) and Monocle (Qiu et al., 2017a; Qiu et al., 2017b; Trapnell et al., 2014).

RNA velocity compares unspliced and spliced transcripts in the single cell dataset, to evaluate the developmental direction of single cells and to generate a UMAP displaying the link between cells (La Manno et al., 2018). Following this, the cluster identities were appended to the RNA velocity map (**Figure 4C**). The map suggests that PL-prolif/PL-Inos are at the origin of most clusters, that PL-0, PL-1, PL-2 and PL-3 are derived from PL-Inos, and that PL-vir1, PL-Rel and PL-robo2 are issued from PL-3, PL-0 and PL-2 (**Figure 4C**). The comparison of the RNA velocity results with the regulons (**Figure 3C**) suggests the pathways involved in the acquisition of the specific properties. First, from PL-prolif to PL-Inos and then to PL-1/PL-3, we observe a gradual reduction of the regulons EcR and E2f1. Then, the JNK associated regulons (Jra, Ets21C) and the regulon Rel become progressively more enriched starting from PL-3 to PL-Rel and PL-vir1 clusters. For the PL-robo2 branch, we observe a gradual enrichment of the regulons associated with the GATA factors Srp and Pnr and with the Hedgehog pathway (Ci) from PL-1/PL-3 to PL-2/PL-0 and then to PL-robo2. Concerning the remaining clusters, PL-Lsp is scattered over the clusters PL-0/PL-1/PL-2/PL-3, suggesting that it is also issued from PL-prolif/PL-Inos, but the directionality is unclear. At last, no clear directionality could be drawn for PL-AMP, PL-Pcd, CC and PL-ImpL2, which suggests that their direct progenitors are not detected in our dataset (**Figure 4C**).

The second approach we used was Monocle, which estimates the cell trajectories by first defining the sequences of gene expression changes required to adopt distinct cell states and then by positioning the cells on the trajectories according to their transcriptomes (Qiu et al., 2017a; Qiu et al., 2017b; Trapnell et al., 2014). The Monocle analysis defines trajectories highly similar to the branches observed with RNA velocity (**Figure 4D**). The PL-prolif/PL-Inos clusters are at one extremity, they are followed by PL-1/PL-3, then PL-2/PL-0, with PL-robo2 at the other extremity. PL-Rel is following the PL-3>PL-0 axis and PL-vir1 follows the same direction than PL-Inos>PL-3.

Overall, these two distinct approaches predict that PL-prolif is at the origin of PL-0 to PL-3, PL-vir1, PL-Rel and PL-robo2. In this model, PL-0 to 3 would represent the bulk of the PL that may then specialize into PL-vir1, PL-Rel and PL-robo.

### Clusters’ dynamics upon wasp infestation

In response to wasp infestation, the *Drosophila* larva displays a strong immune reaction involving multiples organs such as the muscle, the fat body, the lymph gland and the hemocytes (Banerjee et al., 2019). A key response is produced by the hemocytes, with the production of the LM, a fraction of whose aggregate around the wasp egg and encapsulate it to prevent its hatching. During this process, the hemocytes present in the resident compartment and in the lymph gland are recruited through cytokine secretion by the circulating hemocytes, the fat body and the muscles (reviewed in (Banerjee et al., 2019; Kim-Jo et al., 2019; Letourneau et al., 2016)). To characterize the diversity of hemocytes generated by this systemic immune response, including the hemocytes released from the lymph gland, we produced single cell RNA seq data on the cells of the hemolymph from infested larvae.

The efficiency of the immune response of *Drosophila* larvae to wasp infestation critically depends on multiple parameters including the intensity of exposure (number of wasps, duration of the infestation), the genetic background of the *Drosophila* larvae as well as the developmental stage and the temperature at which the infestation occurs (personal observations). We hence optimized the infestation protocol to obtain a rather uniform response and developed a protocol of mild infestation that maximize the survival rate of the *Drosophila OregonR* hosts (see **methods, Figure S6A**). Our protocol maximizes the number of larvae containing a single wasp egg (Bazzi et al., 2018). In these conditions, we observe first the production of LM and a decrease of PL, which suggests that the PL trans-differentiate into LM from the resident and circulating pool of hemocytes (24 h after infestation). Following this, the lymph glands histolyse (>90% of lymph gland histolysed 48 h after infestation) and the hemocytes proliferate (**Figure S6B-D**). By 72 h after infestation (which corresponds to the time of hemocyte collection for the sc RNA seq), all lymph glands are histolysed and the hemolymph contains hemocytes (PL, CC and LM) from the two hematopoietic waves (**Figure S7A-F**) (Bazzi et al., 2018). It is important to note that the WI dataset covers only the cells that have not encapsulated the wasp egg.

Compared to the NI dataset, the single cell data on the WI hemolymph displays two novel cell populations (LM-1 and LM-2) expressing the LM markers (**Figure 5A**, **Figure S3B**). The other clusters are already present in the NI dataset. PL-Inos, PL-prolif, PL-vir1 and PL-0 to 3 clusters increase in cell number, the size of the PL-Lsp, and PL-robo2 clusters remains constant, whereas that of the CC, PL-Pcd and PL-Rel clusters decreases after wasp infestation (**Figure 5B**).

**Figure 5:**
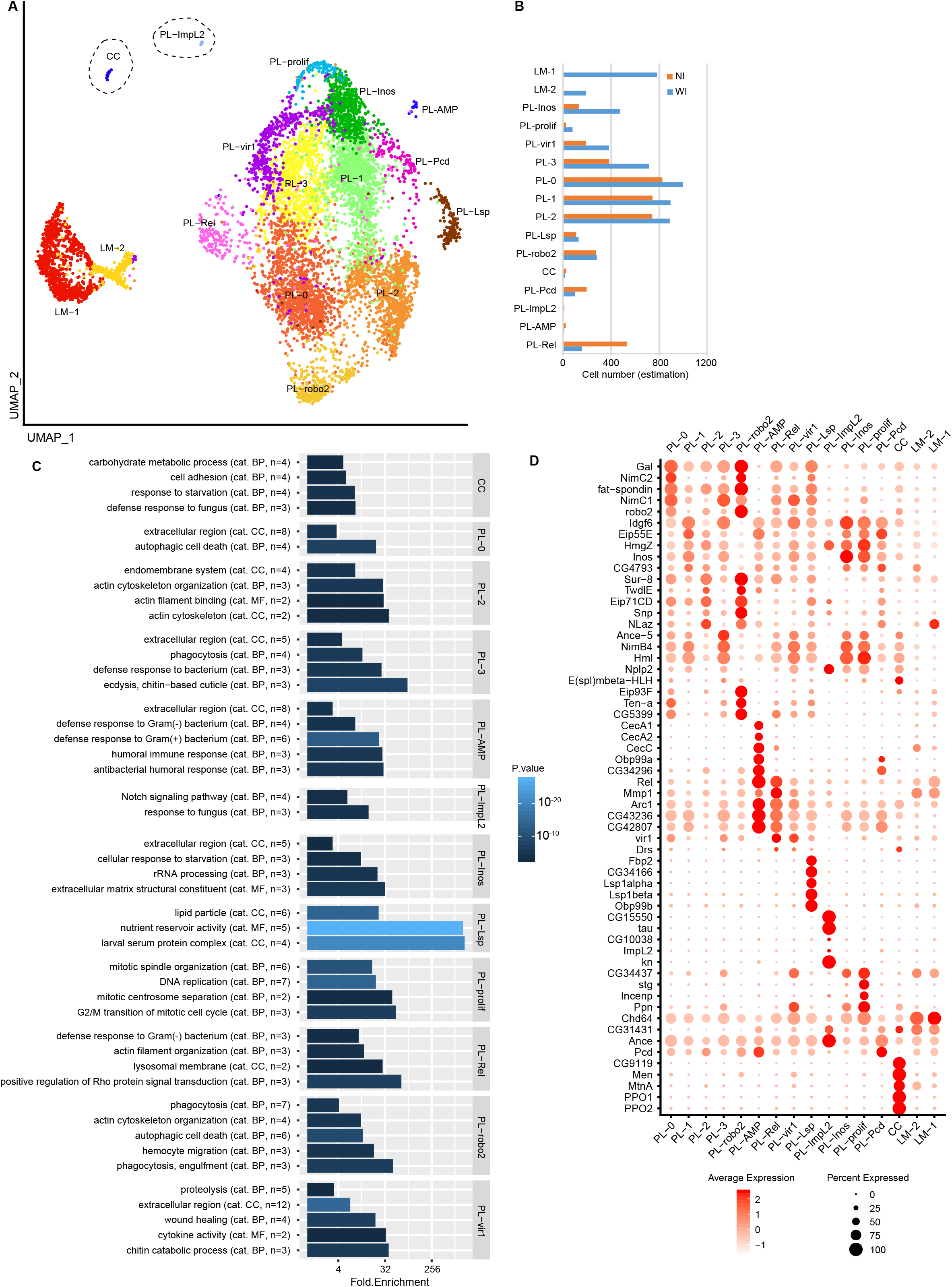
Identification of the hemocyte populations from WL infested by wasp using single cell RNA seq. **A)** UMAP projection representing the 16 clusters of cells identified in the WI sample. Note the presence of two additional clusters, LM-1 and LM-2, compared to the NI sample. The two clusters correspond to the LM. **B)** Bar graph indicating the estimation of the number of cells of each cluster per larva in normal condition (NI, orange) and after wasp infestation (WI, blue). For each cluster, the cell number was deduced from the total number of PL and LM numerated in **Figure S5D**, and the proportion of each cluster in the singe cell datasets. **C)** Bar graph displaying the GO terms enriched in specific clusters for the WI dataset. Represented as in **Figure 2C**. **D)** Dot plot representing the expression levels (gradient of red levels) and the percentage of cells (size of the dots) that express the top 5 markers of each cluster in the WI dataset. Related to **Supplemental Figure S6, S7**.

Comparisons between NI and WI data were carried out for each cluster but did not reveal strong transcriptomic modifications induced by the infestation. Each NI cluster is highly correlated to its WI counterpart **(Figure S6E)** and the markers as well as the GO terms remain similar (**Figure 5C,D**).

### Characterization of the two LM clusters and their developmental links the other clusters

To characterize further the two LM clusters, we compared first their transcriptomes to all the other clusters and carried out GO term enrichment analyses (**Figure 6A,B**). This comparison indicates an enrichment for genes involved in melanotic encapsulation, which is the primary role of the LM, a strong implication of the JNK pathway, which was previously implicated in LM production (Zettervall et al., 2004) and several GO terms related to cytoskeleton reorganization and integrin mediated cell adhesion, two processes that are necessary for the capsule formation around the wasp egg (Irving et al., 2005).

**Figure 6:**
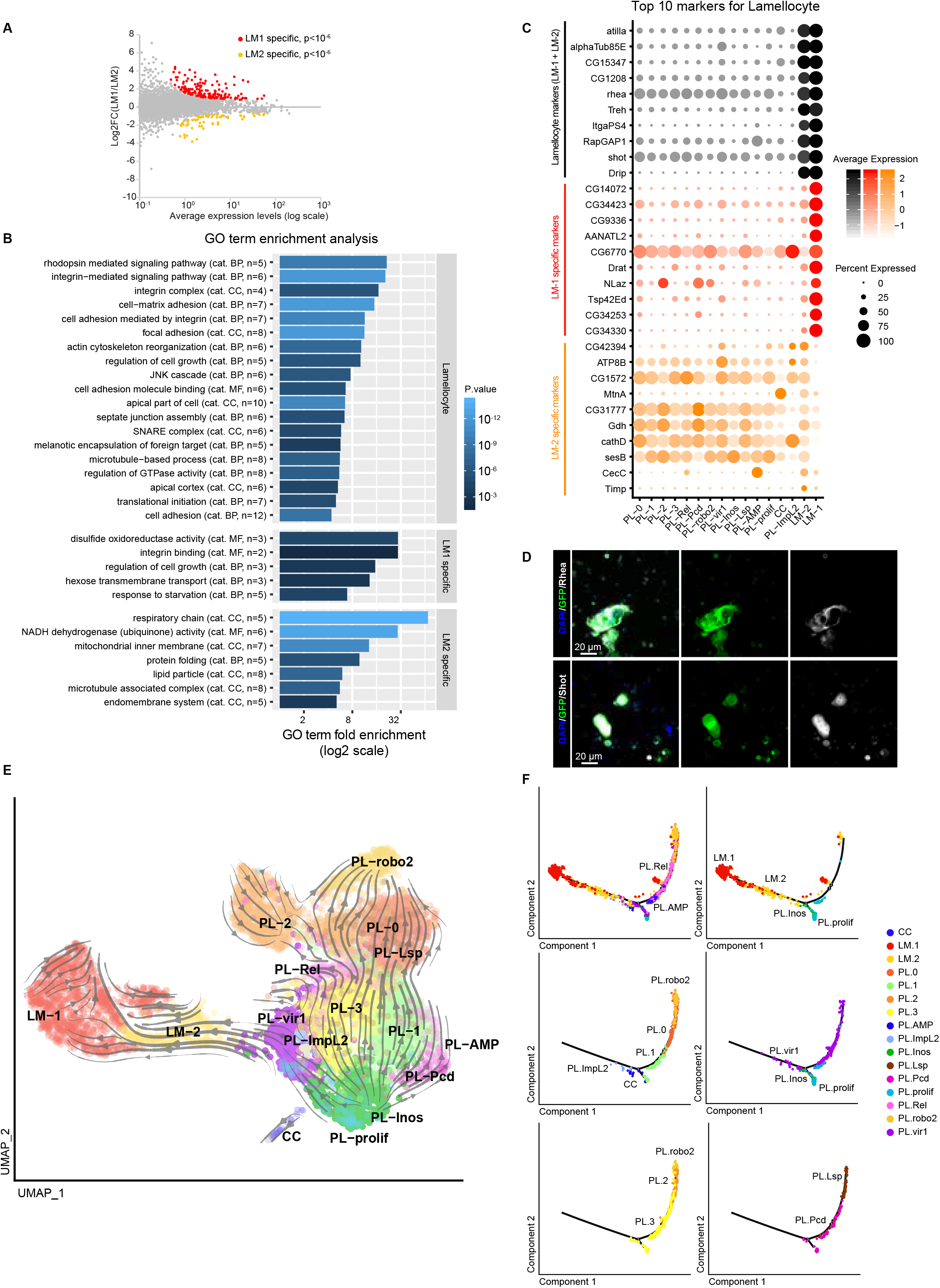
Characterization of the two LM clusters. **A)** Scatter plot comparing the transcriptome of the two LM clusters, deduced from the WI dataset. The x-axis represents the average expression levels of the genes and the y-axis the log2 fold change (LM-1/LM-2). The genes significantly enriched in the LM-1 cluster are highlighted in red (p<10^-6^), those significantly enriched in the LM-2 cluster in orange (p<10^-6^). **B)** GO term enrichment analysis on the genes enriched in LM compared to all other clusters (top panel) and on the genes significantly different in the comparison LM-1 versus LM-2. GO terms enriched in the LM-1 specific genes are in the middle panel and GO terms enriched in the LM-2 specific genes in the lower panel. The bars represent the fold enrichment and the color gradient indicates the p-value of the GO term enrichment (as in **Figure 2C**). **C)** Dot plot representing the expression levels (gradient of color) and the percentage of cells (size of the dots) that express the top 10 markers of LM (in black), of markers specific to LM-1 compared to LM-2 (in red) and specific to LM-2 compared to LM-1 (in orange). Note that most markers enriched in LM-1 compared to LM-2 are exclusively expressed in LM-1 while most markers enriched in LM-2 compared to LM-1 are also expressed in the plasmatocyte and CC clusters. **D)** Confocal images of hemocytes from WL *OregonR* infested by wasp. The immunolabeling was done with antibodies targeting the new LM markers identified in this study Rhea (in gray, **top panels**) and Shot (in gray, **lower panels**). Phalloidin-FITC (in green) labels the actin filament particularly abundant in LM (Tokusumi et al., 2009) and the nuclei were marked with DAPI (blue). Full stacks are displayed, the left panels show the overlay of DAPI, FITC and the LM markers, the middle panels show the FITC alone and the right panels the LM markers alone. The scale bars represent 20μm. **E)** Lineage prediction for the clusters from the WI sample using RNA velocity, as in **Figure 4C**. **F)** Single cell trajectory reconstructed with monocle 2 on the WI dataset. The panels show restricted number of clusters for which the RNA velocity analysis suggested a filiation.

We then compared the LM-1 and LM-2 clusters and found 157 genes up-regulated in LM-1 and 58 genes up-regulated in LM-2 (**Figure 6A**). Only few GO terms are enriched specifically in one or the other cluster: LM-1 specific genes are involved in integrin processing, while LM-2 specific genes are involved in cytoskeleton and mitochondrial processes (**Figure 6B**). The analysis of the top 10 markers for each population is more informative. The markers common between LM-1 and LM-2 include known and novel LM specific markers (including Atilla, ItgaPS4, Rhea, Shot) (in black in **Figure 6C,D**). The markers enriched in LM-1 compared to LM-2 display strong specificity with low expression in LM-2 and in the other hemocyte clusters, whereas the markers enriched in LM-2 are also expressed in most hemocytes clusters (**Figure 6C**). This suggests that LM-1 is constituted by mature LM and LM-2 by cells at a PL/LM intermediate state.

To investigate the link between the two LM populations and the PL, we carried out RNA Velocity and Monocle analyses. The two analyses place LM-2 as intermediate between PL and LM-1, and also suggest that the main cluster producing the LM is PL-vir1. Otherwise, the same major branches from PL-prolif/PL-Inos to PL-robo2, PL-Rel and PL-vir1 as those identified in the NI larvae are observed (**Figure 6E,F**).

### Markers’ dynamics upon WI

Considering the strong immune response induced by the WI, we expected a strong modification of the transcriptional landscape of most hemocytes, however, the transcriptome of the PL clusters remains highly similar in WI compared to NI (see above and **Figure S6E**). To rule out that this observation is due to the low depth of the sc RNA seq, we used qPCR to quantify the main clusters’ markers in NI and WI hemocytes from WL (**Figure 7A**). In agreement with the sc RNA seq data, the large majority of the PL markers maintain the same expression profile upon WI compared to NI. As expected, the LM markers are strongly up-regulated in WI (**Figure 7A**). In addition to the LM markers, the following markers display significant up-regulation: NimB4 (marker of PL-Inos), CecC (PL-AMP), Hml and Sr-C1 (PL-prolif) and Lz (CC). This increase can reflect a modification of the size of their affiliated clusters, which is the case for PL-Inos and PL-prolif (**Figure 5B**). It can also reflect a molecular response to the WI, which is likely the case for CC and PL-AMP whose size remain low upon WI.

**Figure 7:**
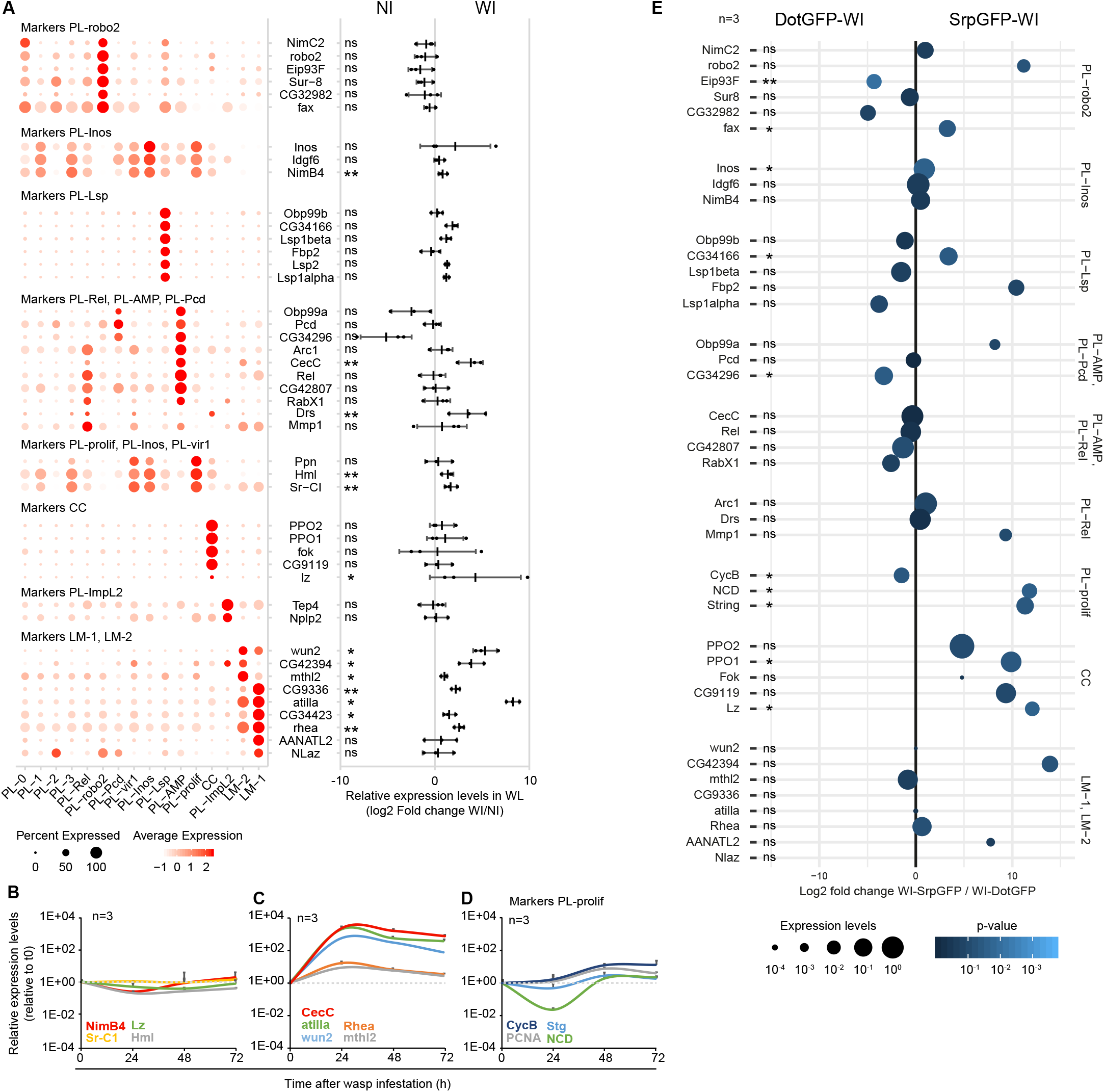
Timing of the production of the different clusters in the hemolymph of WI larvae. **A)** Expression levels of the cluster markers in the hemolymph of WI larvae compared to NI larvae. The dot plot on the left panel indicates the distribution of each marker across all clusters (as in **Figure 2D**), the graph on the right panel, indicates the Log2 of the ratio between the expression level in WI versus NI. Positive values indicate an up-regulation upon wasp infestation and negative values a down-regulation. The experiment was carried out on 3 independent replicates (pool of 10 larvae per replicate). The p-value are estimated by bilateral paired student test and indicated as follow: ns = not-significant (>0.05), “*” = p[0.05 – 0.01[, “**” = p[0.01-0.001[, “***” = p<0.001. **B,C)** Expression levels of the cluster markers increasing in WI compared to NI according to (**A**) during the progression of the immune response to wasp infestation. Collections were carried out at t0 (time of infestation, L2), 24 h (early L3), 48 h (mid L3) and 72 h (WL) after wasp infestation on 3 independent replicates. The x-axis represents the time line and the y-axis the normalized expression levels. The expression levels were normalized to the levels at t0. The markers that do not display strong variability during the timeline are presented in (**B**) and the ones showing a strong up-regulation are in (**C**). **D)** Expression levels during the infestation timeline as in **(B,C)** for the markers of proliferation. **E)** Weighted dot plot representing the enrichment (Log2 fold change) of the markers in hemocytes originating from the lymph gland (DotGFP-WI) compared to embryonic hemocytes (srpGFP-WI) from WI larva. The lymph gland hemocytes and embryonic hemocytes were traced using the lymph gland driver DotGal4 and embryonic driver Srp(hemo)Gal4 respectively and lineage tracing construct and sorted by FACS before quantifying the expression levels by qPCR (**see the method section**). The relative expression levels are indicated by the size of the dots and the pvalue of the log2 fold change with the gradient dark blue (not significant) to light blue (p<10^-3^). The exact p-values are available in Table S6. Related to **Supplemental Figure S6, S8, Table S6.**

To characterize the progression of the immune response to the infestation, we collected the hemocytes 24 h, 48 h and 72 h after WI and quantified the expression levels of specific markers by qPCR (**Figure 7B-D**). CecC and the LM markers undergo strong up-regulation in the 24 h following the WI. Within this timeframe, the lymph gland is still intact (**Figure S6C**) indicating that this early response is produced by the hemocytes of embryonic origin. The high expression levels of the markers are then maintained until 72 h, suggesting that they are also expressed by the hemocytes released from the lymph gland (**Figure 7C**). The other markers (*i.e*. NimB4, Sr-C1, Lz and Hml) do not display strong modulation during the response (**Figure 7B**). We also characterized the expression levels of the PL-prolif markers CycB, PCNA, Stg, and NCD. The 4 markers display stable expression at 24 h and up-regulation at 48 h, which is maintained at 72 h after WI (**Figure 7D**). This suggests that the number of dividing cells stays constant at 24 h and increase later on. These data are concordant with the hemocyte counts at the different stages of infestation (**Figure S6D**). Within 24 h of infestation, the number of hemocytes remains the same but PL transdifferentiate into LM. In the following time points we observe an increase in PL number, which can be explained by the combination of the release of hemocytes from the lymph gland as well as the increased proliferation (**Figure 7D**, **S6D**).

Finally, to determine the origin of the hemocytes, we analyzed the expression levels of the main clusters’ markers in the hemocytes originating from the lymph gland or from the embryo in WI conditions (**Figure 7E**). The hemocytes were traced using the lymph gland specific driver *DotGal4* or the embryonic hemocytes driver *Srp(hemo)Gal4* combined with two lineage tracing transgenes (gtrace, see method section and **Figure S7B,D,F**). WI conditions at WL stage were used to obtain hemocytes from both origins and the cells were sorted by FACS before quantification by qPCR. Of note, the filtration steps necessary for the FACS sorting removed most LM from the samples, which explains the low levels of LM markers in these data (personal observations). The analysis reveals enrichments for the markers of PL-prolif and CC in the embryonic hemocytes. All other clusters display markers in the hemocytes from both origins. This suggests that upon WI, the lymph gland produces hemocytes highly similar to the embryonic hemocytes. It also indicates that the lymph gland releases only few crystal cells and proliferating cells in the hemolymph after WI.

Overall, this analysis indicates that the hemocytes from the lymph gland express most markers found in embryonic hemocytes (in the NI dataset). Thus, at the present level of resolution of the sc RNA seq data, we cannot identify markers specific to the origin (i.e. embryonic or lymph gland) of hemocytes.

### Metabolic properties of the clusters

To observe potential metabolic differences among the clusters, we analyzed the expression profile of the main actors of energy metabolism across the NI and WI datasets (**Figure S8**) in same fashion than on the bulk transcriptomes (**Figure S2**). Although the analyses carried out on the single cell are not as precise as the ones on the bulk transcriptome, most plasmatocyte clusters display metabolic markers in line with the observations made with the bulk RNA seq. We can hence use these data to draw first conclusions.

In the larva, the hemocytes import and metabolize lipids to drive the TCA cycle. Lipid scavenging receptors and the genes of the fatty acid degradation pathway are expressed in most of them. The three exceptions are PL-AMP, PL-Pcd and PL-ImpL2 that express low levels of lipid scavenging receptors and fatty acid degradation genes as well as low levels of TCA genes, suggesting a lower metabolism for these clusters (**Figure S8, left panel**).

The CC distinguish themselves from the PL by the expression of the glucose transporters (Sut1 and Glut1) and genes involved in gluconeogenesis (fbp, Ald) and lipid biosynthesis (Lpin), suggesting a metabolism involving glucose and lipid uptake (**Figure S8, left panel**).

The wasp infestation enhances the expression levels of the lipid scavenging receptors and of the fatty acid degradation genes in the PL, including in the clusters PL-Pcd and PL-AMP, which suggests a stronger metabolic activity (**Figure S8 right panel**). We also noticed increased levels of the glucose transporter Glut1 and of the genes involved in glycolysis in PL-robo2 hemocytes, suggesting that the infestation induces a diversification of the energy source for this cluster (**Figure S8 right panel**). Finally, the LM clusters display a strong expression of the glucose transporter sut1. In addition, LM-1 displays a prominent glycolysis pathway, which may indicate that LM metabolism relies mostly on glucose (**Figure S8 right panel**).

The wasp infestation induces a strong metabolic shift in the larvae where the resources are deviated from development toward the immune response (Rauw, 2012). This shift, mediated by the hemocytes through the production of extracellular adenosine, increases considerably the sugar levels in the hemolymph (Bajgar et al., 2015). Our data suggest that this additional sugar is mostly used by the LM and the PL-robo2 clusters to respond to the wasp infestation, while the other hemocyte clusters keep using lipids.

## Discussion

The immune cells provide the first line of defense against the non-self and accumulating evidence strongly suggest that their function exceeds the immune response. Due to their ability to communicate with the other organs and tissues, immune cells provide ideal sensors for the internal state and homeostasis during development and ontogeny. This raises the issue of immune cell heterogeneity, that is, do different subpopulations display specific potentials? The understanding of immune cell biology heavily relies on the thorough characterization of these cells as well as on the identification of specific markers and subpopulations. This work provides the first atlas of the *Drosophila* hemocytes, by specifically focusing on those that originate from the first hematopoietic wave. We show that these hemocytes undergo a molecular and metabolic shift during development. We show the existence of distinct hemocyte populations and identify a large panel of novel markers specific to the different populations. Monitoring the larval response against the wasp *L. boulardi* reveals the hemocyte behavior upon challenge and defines intermediate and mature LM populations. Finally, we use multiple bioinformatics tools to predict a temporal progression among the different hemocyte clusters in control and in challenged conditions.

### The developmental shift between the embryonic and the larval stages

Immune cells are considered as static components of our defense system, however these cells constantly interact with the ever-changing environment. In addition, the cells that are born in the early embryo experience the extensive rearrangements that occur during development, including tissue and organ formation. We here show that the *Drosophila* hemocytes undergo a significant transcriptional shift that fully complies with the requirements of the embryonic and larval stages.

The highly migratory hemocytes present in the differentiated embryo display a strong developmental role: they allow tissue reshaping by secreting several constituents of the extracellular matrix and by engulfing dead cells through specific scavenger receptors such as NimC4. They also display high levels of gluconeogenesis and TAG synthesis, processes that provide adequate levels of glucose and fatty acid for tissue/organ development. The larval hemocytes, on the other hand, express high levels of transcripts that are linked to the immune response, in accordance with the exposure to pathogens, and are highly phagocytic. Moreover, they express fewer molecules associated with the extracellular matrix as compared to those observed in the embryo. Finally, they strongly express the molecular pathways that release stored energy (beta oxidation, TCA cycle), most likely in preparation for the metamorphosis and to help building the adult tissues.

### Different types of hemocytes in wild type larvae

The single cell analysis on the NI animals reveals the presence of 14 different hemocyte clusters, in addition to the classical distinction between PL and CC. This provides us with a battery of novel specific markers (**Table S7**) that will make it possible to investigate the role of the different hemocyte populations and to generate more targeted genetic tools. Excitingly, we can already define distinct features and functions of the different clusters, based on the profile of gene expression, on the enrichment in specific GO terms and regulons as well as on the *in vivo* validation. Indeed a number of clusters is identified by a single regulon (Tbp for PL-Lsp, lz, for CC, E2f1 for PL-prolif) or by a specific combination of regulons in the case of related clusters (e.g. PL-vir1, PL-Rel and PL-Amp).

The **PL-Rel** cluster (12%) likely provides a cellular reservoir for a specific immune response, the closely related **PL-AMP** hemocytes (0.5%) seem more specifically dedicated to the humoral response, whereas **PL-vir1** hemocytes (4%) seem dedicated to the anti-viral response. These three clusters share GO terms and regulons associated with immune functions, suggesting that they respond to a variety of challenges.

The **PL-Lsp** hemocytes (<3%) represent the nutrient reservoir that stores amino acids and, in agreement with a role in homeostasis, they are mostly located in circulation. The PL-Lsp and PL-AMP hemocytes are associated with the major roles of the fat body, the metabolic homeostasis and the humoral immune response, suggesting that they contribute to the fat body – hemocyte axis acting in physiological and pathological conditions. This axis is bidirectional. For example: 1) the small secreted peptide Edin produced in the fat body controls the number of plasmatocytes in circulation upon wasp infestation; 2) the hemocyte expression of the Spaetzle ligand controls the activity of Toll signaling in the fat body and affects the response to infection (Shia et al., 2009) as well as tumor growth (Parisi et al., 2014); 3) the metabolically induced production of the NimB5 protein from the fat body adjusts the number of hemocytes to the physiological state of the larva (Ramond et al., 2019).

The **PL-robo2** hemocytes (6% of the total population) are associated with phagocytosis but are also enriched for the lipid scavenger receptor Crq involved in the inflammatory response upon high fat diet (Woodcock et al., 2015). This cluster shares features with the large **PL-0** and **PL-2** clusters that are mildly enriched for the regulon related to the phagocytic abilities (srp), in agreement with the finding that the vast majority of the larval hemocytes is phagocytic (**Figure 1**). The **PL-0, PL-2, PL-1** and **PL-3** large clusters, which do not display strong specific molecular features, may indeed represent cells that can serve different purposes, perhaps less efficiently than the more specialized hemocytes. They may also constitute a reservoir of cells that have a basal activity but express enhanced potentials in response to challenges, in line with the trajectories identified by the bioinformatic analyses.

The data on the small **CC** cluster validate the role of these cells in melanization and reveal a distinct metabolism compared to PL. The CC seem to use glucose as energy source and the PL mostly lipids.

The genes associated with mitosis are enriched in a single cluster that preferentially localizes to the resident compartment. This indicates that **PL-prolif,** which represents less than 1% of the total population, provides the pool of mitotic precursors in the larva. In agreement with these data, the bioinformatics analyses reveal a sequential progression leading from PL-prolif to most of the hemocyte clusters identified in the larva. **PL-Inos**, which is the cluster the most closely related to PL-prolif, is also associated with the resident compartment and likely represents immature progenitors.

According to the bioinformatics analyses, the **PL-Impl2** hemocytes seem not to originate from PL-prolif hemocyte. These may represent hemocytes that are set aside in the embryo and indeed the comparison of all the clusters reveals that the PL-Impl2 hemocytes are significantly enriched for transcripts that are specific to the E16 hemocytes (bulk transcriptome analyses, personal communication).

The identification of different populations of specialized plasmatocytes in the *Drosophila* larva prompts us to draw parallels with the mammalian immune cells. The closest relative to plasmatocytes are the monocytes and the macrophages (Wood and Martin, 2017). Monocytes are equipped with Toll-like receptors, scavenger receptors and their main function is to patrol as well as remove microorganisms, lipids and dying cells via phagocytosis. Upon inflammation, they infiltrate specific tissues and differentiate into macrophages. The macrophages keep phagocytosing, induce an inflammatory response by releasing cytokines and participate to the repair of the tissue (Yang et al., 2014). The sc RNA seq assay reveals that hemocytes express markers such as Integrin alphaPS2 (If), EcR/Hr96, Lamp1, Rgh, Tfc/lectin-46Cb/CG34033 and Lz, which are the *Drosophila* orthologues of CD11b, PPARγ, CD68, Dectin, CD207 and RUNX, respectively. In mammals these proteins are responsible for migration, adhesion, phagocytosis, differentiation from monocytes to macrophages and pathogen recognition (Daley et al., 2017; Heming et al., 2018; Podolnikova et al., 2016; Ramprasad et al., 1996; Voon et al., 2015).

### The larval response to wasp infestation

The single cell analysis upon WI reveals the reduced representation of some clusters such as PL-Rel and the expanded representation of ‘early’ clusters (e.g. PL-prolif, PL-Inos). Thus, specific hemocyte clusters may preferentially survive/proliferate upon challenge. The majority of the clusters, however, remain equally represented in the two conditions and the correlation between the average transcriptomes in NI and WI conditions reveals strong similarity between most of the identified clusters (Pearson = 0.97). This implies that the hemocytes produced by the 1^st^ and the 2^nd^ hematopoietic waves share major features.

Two new populations of cells appear, LM1 and LM2, the second one representing an intermediate state characterized by the co-expression of LM and PL genes. Interestingly, LM2 also expresses a specific identity that is linked to energy supply (e.g. respiratory chain, NADH activity), whereas LM1 cells are mostly devoted to encapsulation.

According to the bioinformatics predictions (RNA Velocity and Monocle) on the WI dataset, the PL-prolif cluster seems to rapidly branch out: one arm gives rise to the LM clusters, partly associated with the PL-vir1 cluster, whereas the other gives rise to the other plasmatocyte clusters. This early separation could support the hypothesis of a dedicated precursor, the lamelloblast, producing a second population of LM that is not generated through plasmatocyte trans-differentiation (Anderl et al., 2016). In this model, the 1^st^ hematopoietic wave would produce LM through trans-differentiation (the expression of LM2 markers only increases in the first 24 h after infestation), whereas the 2^nd^ wave would do it (also) through the mitotically active lamelloblast. At the level of resolution provided by the sc RNA seq assay, we may have lost the lamelloblast cluster.

In sum, the different clusters identified by the sc RNA seq assay exhibit distinct features, which can be now tested functionally using the newly identified markers and the associated genetic tools that are publically available (Gal4 drivers, RNAi and overexpressing transgenes, mutations). Future technological refinements may enhance the depth of the analyses, as the current sc RNA seq assays only allows for the identification of a subset of genes for each cluster, the most expressed ones. As an example, the larval hemocytes do not all phagocytose with the same efficiency, but we cannot allocate the different potentials to specific clusters (**Figure S5F**). Nevertheless, our data on the bulk and single cell transcriptomes of the *Drosophila* hemocytes provide a powerful framework to understand the role of immune cells in physiological and pathological conditions.

## Acknowledgments

We thank the Imaging Center of the IGBMC for technical assistance. This study was supported by the grant ANR-10-LABX-0030-INRT, a French State fund managed by the Agence Nationale de la Recherche under the frame program Investissements d’Avenir ANR-10-IDEX-0002-02. The sequencing was performed by the GenomEast platform, a member of the ‘France Génomique’ consortium (ANR-10-INBS-0009). We thank I. Ando, K. Bruckner, M. Crozatier, M. Meister, D. Siekhaus, D. Bohmann for providing fly and wasp stocks. In addition, stocks obtained from the Bloomington Drosophila Stock Center (NIH P40OD018537) as well as antibodies obtained from the Developmental Studies Hybridoma Bank created by the NICHD of the NIH and maintained at The University of Iowa (Department of Biology, Iowa City, IA 52242) were used in this study. We thank Dasaradhi Palakodeti for useful discussion on transcriptome analysis.

This work was supported by INSERM, CNRS, UDS, Ligue Régionale contre le Cancer, Hôpital de Strasbourg, ARC, CEFIPRA, ANR grants and by the CNRS/University LIA Calim. T. Mukherjee and Nivedita Hariharan are funded by the Council for Scientific and Industrial Research fellowship. The IGBMC was also supported by a French state fund through the ANR labex.

## Author contributions

Conceptualization, P.B.C. and A.G.; Methodology, P.B.C., A.G., R.S., A.P., C.D., N.M., A.R., T.M., N.H.; Investigation, P.B.C., A.G., R.S., A.P., C.D., N.M., A.R., T.M., N.H.; Writing–Original Draft, P.B.C., A.G., R.S., A.P.; Writing–Review & Editing, P.B.C., A.G., R.S., A.P.; Funding Acquisition, A.G., N.M., T.M.; Resources, A.G.; Supervision, P.B.C. and A.G..

## STAR Methods

### Fly strains and genetics

All flies were raised on standard media at 25°C. For the bulk sequencing, the hemocytes from stage 16 (E16) embryos were collected from *srp(hemoGal4/+;UAS-RFP/+* animals obtained upon crossing *srp(hemo)Gal4* (gift from K. Brückner) (Bruckner et al., 2004) and *UAS-RFP* flies (RRID:BDSC_8547). The wandering L3 (WL) hemocytes were collected from staged *HmlΔRFP/+* animals upon crossing *HmlΔRFP* (Makhijani et al., 2011) with *OregonR* flies (108-117 h After Egg Laying, h AEL).

For the single cell sequencing, *OregonR*-flies were used as the wildtype (WT) strain for all the experiments and for the single cell RNA sequencing.

Validation of the single cell data involved the following stocks: *srp(hemo)-moesin-RFP* (stock D2244 on chr 2, gift from D. Siekhaus (Gyoergy et al., 2018)), *Mimic-Lsp1beta-MI05460* (RRID:BDSC_40782), *Lsp2-Gal4* (RRID:BDSC_6357), the lineage tracing line *UAS-FLP,Ubi-p63E(FRT.STOP)Stinger* (RRID:BDSC_28282) that was combined with *UAS-FLP; act5c-FRT,y+,FRT-Gal4,UASmCD8GFP* (gift from I. Ando (Honti et al., 2010)), *Dot-Gal4* (RRID:BDSC _6903), *GstD-LacZ* (gift from D. Bohmann (Sykiotis and Bohmann, 2008)).

### FACS sorting of embryonic and larval hemocytes

Staged egg laying was carried out to produce E16 embryos as follows. The cross to produce *srp(hemo)Gal4/+;UAS-RFP/+* embryos (with at least 100 females) was transferred to egg laying cages on a yeasted apple juice agar at 25°C. After a pre-lay period of 30 min, the agar plates containing yeast were replaced with fresh plates and flies were left to lay for 3 h at 25°C. Agar plates were then removed and the embryos were raised for 11 h and 40 min at 25°C until they reached stage 16. Embryos were then isolated from the medium and washed on a 100 μm mesh. The collected embryos were transferred into a cold solution of phosphate-buffered saline (PBS) in a Dounce homogenizer on ice. The embryos were dissociated using the large clearance pestle then the small clearance pestle and then the cells were filtered through a 70 μm filter to prepare them for FACS sorting. The cells were sorted using FACS Aria II (BD Biosciences) at 4°C in three independent biological replicates. Live cells were first selected based on the forward scatter and side scatter and only single cells were taken into account. *OregonR* cells were used as a negative control to set the gate for the sorting of RFP positive cells only (**Figure S1**). RFP positive hemocytes were collected in 1 ml of TRI reagent (MRC) for RNA extraction.

For the wandering L3 *HmlΔRFP/+* hemocytes, staged lay of 3 h were carried out at 25°C to prevent overcrowding of the vials (between 50 and 100 embryos per vial) and wandering larvae were collected 108-117 h AEL, bled in cold PBS containing PTU (Sigma-Aldrich P7629) to prevent hemocyte melanization (Lerner and Fitzpatrick, 1950), filtered through a 70 μm filter to isolate them and sorted by FACS as described for the embryonic hemocytes.

The purity of the sorted populations was assessed prior to the collection of the sample for RNA extraction by carrying out a post–sort step. The FACS sorter was set up to produce hemocyte pools displaying at least 80% of purity on the post-sort analysis.

### RNA extraction and bulk RNA sequencing

The sorted cells were homogenized then left at room temperature (RT) for 5 min to ensure complete dissociation of nucleoprotein complexes. 0.2 ml of chloroform was added to each sample followed by centrifugation at 12,000g for 15 min at 4°C. The upper aqueous phase containing the RNA was collected and transferred to a fresh autoclaved tube. 0.5 ml of 2-propanol were added, and the samples were incubated for 5–10 min at RT. The RNA was precipitated by centrifugation then washed with 1 ml of 75% ethanol then precipitated again and air dried. 20 μl of RNase-free water was added to each sample before incubation at 55°C for 15 min. Single-end mRNA-Seq libraries were prepared using the SMARTer (Takara) Low input RNA kit for Illumina sequencing. All samples were sequenced in 50-length Single-Read. At least 40.10^6^ reads were produced for each replicate (**Figure S1**).

### Analysis of bulk RNA seq data

The data analysis was performed using the GalaxEast platform, the Galaxy instance of east of France (http://www.galaxeast.fr/, RRID:SCR_006281) (Afgan et al., 2018). First, summary statistics was computed on the raw FastQ Illumina files of the dataset using the quality control tool for high throughput sequence data FastQC (Babraham Bioinformatics, RRID:SCR_014583). FastQ Illumina files were converted to FastQ Sanger using the FastQ Groomer tool after assessing the quality of the sequencing. The FastQ Sanger files were then mapped onto the *Drosophila melanogaster* reference genome Dm6 using TopHat (RRID:SCR_013035) (Trapnell et al., 2009). As for the expression levels, the analysis of differential gene expression was based on the number of reads per annotated gene. This was done by using Htseq-Count (RRID:SCR_011867) (Anders et al., 2015) and the comparison and normalization of the data between the different cell types was done in Deseq2 (**Figure S1)** (RRID:SCR_015687) (Anders and Huber, 2010). The gene ontology studies presented in **Figure 1B** and **Supplemental Table S1** were done using the Database for Annotation, Visualization and Integrated Discovery (DAVID) v6.8 (https://david.ncifcrf.gov/, RRID:SCR_001881) (Huang da et al., 2009) for the identification of biological processes.

The metabolic pathway analysis (**Figure S2**) was done as follow. Genes that showed a fold change ≥ 2 with an adjusted p-value of less than 0.05, were considered for gene set enrichment analysis. Gene ontology and KEGG pathway enrichment analysis of the differentially expressed genes was done in ShinyGO v0.60 webserver (Ge and Jung, 2018). Genes associated with metabolic pathways considered in this study, were retrieved from the KEGG database (http://www.genome.jp/, RRID:SCR_012773) (Kanehisa and Goto, 2000). The log2FC values of the metabolic genes (q<0.05) in the hemocytes, was then plotted using R (version 3.4.0) (R Core Team, 2017). The corresponding expression data for these genes in the ‘Embryo_16-18hr’ and ‘larva_L3_puffstage_7-9’ developmental stages from modEncode database (RRID:SCR_006206) (Graveley et al., 2011), was downloaded using the webtool DGET(Hu et al., 2017) (https://www.flyrnai.org/tools/dget/web/). These data were used to calculate the fold change and is represented as Bar-plots. For genes with paralogs, the paralog with highest fold-change has been considered for the analysis. The details of all the genes (including all the paralogs) from these pathways are present in **Table S6** and the genes represented in the bar-plots (**Figure S2**) are highlighted in yellow.

### Phagocytosis Assay

Hemocytes from E16 embryos and wandering larvae underwent the phagocytosis assay with latex beads. Briefly, *srp(hemo)-moesin-RFP* flies were staged for 3 h at 25°C and then the embryos were incubated at 25°C for 12 h in order to reach stage 16. Then the embryos were collected and dechorionated in 25% bleach for 5 min at RT. Upon that the embryos were washed, homogenised with Dounce Homogenizer in Schneider medium complemented with 10% Fetal Calf Serum (FCS), 0.5% penicillin, 0.5% streptomycin (PS), and few crystals of N-phenylthiourea ≥98% (PTU) (Sigma-Aldrich P7629) to prevent hemocyte melanization (Lerner and Fitzpatrick, 1950) and filtered with a 70 μm filter. 20 third instar larvae were bled in Schneider medium. Hemocytes for both stages were treated at the same time with latex beads 0.50 μm (Polysciences Inc., cat 17152) for 5 and 20 min, cytospinned at 700 rpm for 3 min, fixed for 10 min in 4% paraformaldehyde/PBS at RT, incubated for 30 min with DAPI to label nuclei (Sigma-Aldrich) (diluted to 10-3 g/L in blocking reagent) and phalloidin Cy3 (only for the WL3 hemocytes due to low moesin RFP signal), and then mounted in Aqua-Poly/Mount (Polysciences, Inc.). The slides were analyzed by confocal microscopy (Leica Spinning Disk) using identical settings.

For the phagocytosis assay on NimC1/P1 negative hemocytes, 20 third instar larvae were bled in Schneider medium supplemented with PTU and were treated with latex beads 0.50 μm diluted 1/500 for 2, 5 or 10 min. Cells were then fixed and labelled with Rabbit anti-Srp(Bazzi et al., 2018), mouse anti-P1(Vilmos et al., 2004). We then used the secondary antibodies Cy5 Goat Anti-Mouse IgG (Jackson ImmunoResearch Labs Cat# 115-177-003, RRID:AB_2338719) and Cy3 Goat Anti-Rabbit IgG (Jackson ImmunoResearch Labs Cat# 111-165-144, RRID:AB_2338006) and DAPI. Images were acquired using Leica Spinning Disk microscope. Images produced were analysed in Fiji (RRID:SCR_002285) (Schindelin et al., 2012).

### Imaging

The images produced for this paper were acquired on a Leica Spinning Disk from the Imaging center of the IGBMC (http://ici.igbmc.fr/). The acquisition step was 0.5 μm with a 40X magnification. For the quantifications, three or more fields per sample were used with more than 50 cells in total. The intensity of latex beads or protein levels were measured with the Imaris software (version 9.5).

### Hemocyte immunolabeling

10 3^rd^ instar larvae per sample were bled in Schneider medium complemented with 10% Fetal Calf Serum (FCS), 0.5% penicillin, 0.5% streptomycin (PS), and few crystals of N-phenylthiourea ≥98% (PTU). For the collection of circulating hemocytes, the hemolymph was gently allowed to exit, while resident hemocytes were scraped and/or jabbed off the carcass in a second well as described in (Petraki et al., 2015). For the infested larvae, there was no separation of circulating from resident hemocytes and we used 5 larvae per sample. The cells were cytospinned at 700 rpm for 3 min, then the samples were fixed for 10 min in 4% paraformaldehyde/PBS at RT, incubated with blocking reagent (Roche) for 1 h at RT, incubated overnight at 4°C with primary antibodies diluted in blocking reagent, washed three times for 10 min with PTX (PBS, 0.1% triton-x100), incubated for 1 h with secondary antibodies, washed twice for 10 min with PTX, incubated for 30 min with DAPI and phalloidin GFP, and then mounted Aqua-Poly/Mount (Polysciences, Inc.). The slides were analyzed by confocal microscopy (Leica Spinning Disk) using identical settings between control and infested samples. The following combination of primary antibodies was used to determine the fraction of LM: mouse anti-Relish [1:40; supernatant from the Developmental Studies Hybridoma Bank (DSHB Cat# anti-Relish-C 21F3, RRID:AB_1553772)], mouse anti-Shot (1:40; supernatant, DSHB Cat# anti-Shot mAbRod1, RRID:AB_528467), mouse anti-Talin (Rhea) (1:40; DSHB Cat# Talin A22A, RRID:AB_10660289) mouse anti-Talin (Rhea) (1:40,DSHB Cat# Talin E16B, RRID:AB_10683995) and rat anti-Elav (1:200; DSHB Cat# Rat-Elav-7E8A10 anti-elav, RRID:AB_528218). The secondary antibodies, Cy3 Donkey Anti-Mouse IgG (Jackson ImmunoResearch Labs Cat# 715-165-151, RRID:AB_2315777), Cy3 Goat Anti-Rat IgG (Jackson ImmunoResearch Labs Cat# 112-165-167, RRID:AB_2338251), Cy5 Goat Anti-Mouse IgG (Jackson ImmunoResearch Labs Cat# 115-177-003, RRID:AB_2338719) and Cy5-AffiniPure Goat Anti-Rat IgG (H+L) (Jackson ImmunoResearch Labs Cat# 112-175-167, RRID:AB_2338264) were used at 1:500.

### Quantitative PCR

For the comparison between resident and circulating hemocytes, 20 3^rd^ instar larvae per sample were bled on ice cold PBS and the circulating hemocytes were separated from the resident ones as described above. The cells were then centrifuged at 1200 rpm, 4°C and RNA isolation was performed with the RNeasy mini kit (Qiagen) by following the manufacture’s protocol. The DNase treatment was performed with the TURBO DNA-free kit (Invitrogen) and the reverse transcription (RT) was done by using the Super-Script IV (Invitrogen) with random primers. The cycle program used for the RT 65°C for 10 min, 55°C for 20 min, 80°C for 10 min. The qPCR we used FastStart Essential DNA Green Master (Roche). The primers are listed in **Supplemental Figure S5**. The p-values and statistical test used are indicated in **Supplemental Figure S6.**

For the quantitative PCR done on *UAS-FLP/+;srp(hemo)Gal4/gtrace-mCD8-GFP;gtrace-nls-GFP/+* and *UAS-FLP/+;DotGal4/gtrace-mCD8-GFP;gtrace-nls-GFP/+*, 30 wandering third instar larvae were collected per replicate, the larvae were bled in PBS supplemented with PTU and the cells were filtered through a 70 μm filter to prepare them for FACS sorting. The cells were sorted using FACS Aria II (BD Biosciences) at 4°C in three independent biological replicates. Live cells were first selected based on the forward scatter and side scatter and only single cells were taken into account. *UAS-FLP/+;gtrace-mCD8-GFP/+;gtrace-nls-GFP/+* cells were used as a negative control to set the gate for the sorting of GFP positive cells only. GFP positive hemocytes were collected and the RNA isolation and qPCR were done as mentioned above.

### Single cell sample preparation, sequencing and analysis

*OregonR* females were used for the generation of the single cell data. For the NI sample, 20 female larvae were collected at the wandering L3 stage (108 to 117 h AEL at 25°C) and bled in Schneider medium complemented with PTU on ice. Both circulating and resident pools of hemocytes were collected.

For the WI sample, staged larvae were infested at the L2 stage (48 to 56 h AEL) for 2 h at 20°C with 20 female wasps (*L. boulardi*) per 100 *Drosophila* larvae. Following infestation, the larvae were raised at 25°C until the wandering L3 stage. Of note, the wasp infestation induces a developmental delay, as infested larvae reach the wandering stage at ~120 h AEL. In an effort to obtain reproducible results, the infestation conditions were optimized so that the majority of the larvae carry a single wasp egg (Bazzi et al., 2018) 20 female larvae were processed as described above for the NI condition.

After bleeding, the hemolymph from NI and WI larvae was filtered on a 100 μm mesh to remove cell aggregates and wasp eggs. Cell concentration was estimated with a hemocytometer. 10,000 cells of each condition were used to prepare the 3’mRNA seq libraries with the Chromium Single Cell 3’ Reagent Kits v2 (10x Genomics). The libraries were then sequenced on the sequencer Illumina Hiseq 4000, on two lanes using paired sequencing of 2×100nt. The raw data were analysed using Cell Ranger v3.0.1 (pipeline from 10x Genomics) and mapped to the *Drosophila* genome assembly BDGP6_ens95.

### Clustering the single cell data

The single cell data were further analysed using the R based toolkit Seurat v3 (RRID:SCR_016341) (Butler et al., 2018; Stuart et al., 2018). Both NI and WI datasets were combined using the standard workflow for data integration. The variable features were selected with the variance stabilizing transformation method. The optimal number of dimensions for the generation of the UMAP was determined using the tools DimHeatmap and Elbowplots, the clustering was done on 20 dimensions with a resolution of 0.55. These parameters returned 14 clusters. Further clustering was then attempted on each cluster separately (manual curation). If the manual curation returned more than 10 distinct markers (|avg_logFC| >1, ROC analysis returning AUC > 0.75) the cluster was subdivided. This protocol led to the subdivision of the PL-Rel and PL-Inos clusters (see **Figure S3**).

The GO term enrichment analysis was carried out based on gene expression levels in NI condition to identify the main features of the clusters. The number of LM being negligible in the NI dataset (8 cells), they were excluded from this analysis. The genes enriched in each cluster (log2FC > 0.25, adjusted p-value < 0.01, determined with FindAllMarkers from Seurat, **Supplemental Table S2**) were analysed using DAVID. The list of cluster specific genes was compared to the list of genes expressed in the whole dataset. The whole GO term results are available in **Supplemental Table S3** and for each hemocyte cluster, the hemocyte related GO terms displaying the strongest enrichment are presented in **Figure 2C**.

### RNA velocity analysis

To visualize the ongoing transcriptional changes in single cell, we adopted the approach described in (La Manno et al., 2018), i.e. we calculated the “velocity” of each cell in the highdimensional gene expression space. We started by generating the loom file, containing spliced and unspliced reads with velocyto, version 0.17.17 (http://velocyto.org/velocyto.py) (La Manno et al., 2018). As input to velocyto, we gave the ‘run10x’ option (specific for 10x data), the CellRanger output and Ensembl annotation v95 (ftp://ftp.ensembl.org/pub/release-95/gtf/drosophila_melanogaster/). Our NI dataset includes 4.2% intronic sequences, on which velocyto analysis was based. The representation of the data was done with the package scVelo (https://scvelo.readthedocs.io/) (Bergen et al., 2019) that implements UMAP representations.

### Monocle graph reconstruction

To complement the RNA velocity analysis, we run also Monocle 2 (Qiu et al., 2017a; Qiu et al., 2017b; Trapnell et al., 2014). We use the R version 3.5.1. The CellDataSet has been built using ‘expressionFamily=negbinomial.size()’. We adopted ‘DDRTree’ method for dimensionality reduction and a maximum number of components equals 2.

### Regulon analysis

To identify the regulons involved in the hematopoietic system, we ran Single-Cell regulatory Network Inference and Clustering (SCENIC, RRID:SCR_017247)(Aibar et al., 2017) through its Python implementation pySCENIC, version 0.9.19 (https://pyscenic.readthedocs.io/en/latest/). The source code was downloaded from the GitHub repository https://github.com/aertslab/pySCENIC.git. The supplemental files necessary to run SCENIC were obtained from https://resources-mirror.aertslab.org/cistarget/. For the analysis, we chose the motifs version 8 (cistarget/motif2tf/motifs-v8-nr.flybase-m0.001-o0.0.tbl) and the regulatory elements within 5kb upstream the TSS and the transcript introns (cistarget/databases/drosophila_melanogaster/dm6/flybase_r6.02/mc8nr/gene_based/dm6-5kb-upstream-full-tx-11species.mc8nr.feather). Finally, to identify the most significant regulons showing a different activity among clusters, we performed a Mann-Whitney U test (Mann and Whitney, 1947), between the AUC scores given by SCENIC in a specific cluster versus all the rest of the clusters.

### Comparison of regulon-based and gene-based clusters

We performed clustering based on the regulon AUC scores per cell, by adopting the ‘Louvain’ algorithm with a resolution equal to 1.3 in Scanpy (Wolf et al., 2018) version 1.4.4, in order to obtain the same number of clusters (=16) than in the previous gene-based analysis. To compare the results from gene-based and the regulon-based clustering, we calculated the Rand index(Rand, 1971), obtaining a value of 0.83. Rand index (RI) = 0 means no overlaps between the clusters, RI = 1 means perfect overlap.

## Supplemental information

### Supplemental Figures

**Figure S1**: Outline of the experimental design used to generate the transcriptomes of the embryonic and larval hemocytes.

**A)** Hemocyte suspensions are prepared from stage 16 embryos (E16) *srp(hemo)Gal4/+;UAS-RFP/+* (right panel) and wandering L3 (WL) *HmlΔRFP* (left panel), the left panel shows the endogenous RFP in a living larva and the right panel shows immunolabeling with an anti-RFP antibody and DAPI.

**B)** The embryonic hemocytes are then sorted according to the RFP signal by FACS.

**C)** After RNA extraction, poly-A RNAs are selected to make the libraries. Following the single end sequencing (50nt), the quality of the data is assessed by FastQC, the sequences are aligned with Tophat and compared with HTseq and DESeq2. Three biological replicates are generated per stage. The number of reads per sample is indicated to the right.

**D)** Scatter plots highlighting in black the subset of genes associated with the GO term extracellular matrix. Light gray dots indicate genes that are not expressed in a significantly different manner between E16 and WL. Dark gray dots indicate genes that are expressed in a significantly different manner. Genes found in the *in situ* databases are indicated. Note that these genes are among the ones presenting the highest expression levels.

**E)** Expression profiles of extracellular matrix genes in the embryos produced by *in situ* hybridization. The pictures were retrieved from the two databases BDGP *in situ* database and FlyFish *in situ* database.

**Figure S2: Metabolic pathway enrichment analysis in bulk E16 and WL hemocyte transcriptomes**

**A,B)** Bar chart representing the expression levels (Log2 fold change WL/E16) of rate-limiting metabolic genes differentially expressed in the larval versus embryonic hemocytes. The left panels show the comparison of whole WL to whole E16 embryos (from modEncode) and the right panels the comparison of WL to E16 hemocytes (our data). The role in metabolism is indicated on the left side of the graphs.

**C)** Graphical illustration of metabolic pathways differentially regulated in hemocytes. Green indicates up-regulated genes and red indicates down-regulated genes in larval hemocytes.

**Figure S3: Manual curation of the single cell data (NI + WI)**

**A,A’)** UMAP projections before (**A**) and after (**A’**) manual curation of the clusters. Note the subdivision of cluster 7 (CL 7) into two related clusters, PL-Inos and PL-prolif, as well as that of cluster 4 (CL 4) into the PL-Rel and PL-AMP related clusters. Similar attempts to subdivide the other clusters did not lead to the identification of additional related clusters. Since we did not find major gene expression differences between NI condition and upon Wasp infestation (apart from the appearance of the LM clusters), the name of each cluster was assigned after merging the two datasets.

**B)** Dot plot representing the expression levels (green to red gradient) and the percentage of cells (size of the dots) expressing markers that are known to be specific to the CC, to the LM or to PL (x-axis), for each cluster (y-axis).

**C-C’’,D-D’’)** UMAP projections highlighting the two clusters before (**C,D**) and after subdivision **(C’,D’**) (see manual curation in methods). The expression of the main markers is displayed in the violin plots (**C’’,D’’**). Each violin plot represents the expression levels of a selected gene within each cell of the cluster (black dots) and the cell distribution is highlighted by the shaded area (green for PL-Inos, light blue for PL-prolif, pink for PL-Rel and dark blue for PL-AMP) (**C’’, D’’**). The genes were selected for being either markers in common between the related clusters (Inos for PL-Inos and PL-prolif; Rel for PL-Rel and PL-AMP) or markers allowing the distinction between the two related clusters. Pen, Chd64, PCNA and CycB are significantly more expressed in the PL-prolif cluster. CG8501 and Gal are more expressed in the PL-Rel cluster, whereas CG34296 and CecA1 are more expressed in the PL-Rel-AMP cluster.

**Figure S4: Expression profiles of known hemocytes markers and ‘extracellular region’ genes in the NI dataset**

**A)** Dot plot representing the expression levels (green to red gradient) and the proportion of cells (size of the dot) of known markers of CC and PL across the different hemocyte clusters of the NI dataset.

**B)** Dot plot of genes annotated as ‘extracellular region’ and enriched in specific clusters from the NI dataset.

**Figure S5: Validation of the NI clusters**

**A)** Dot plot of genes enriched in the PL-Lsp cluster in the NI dataset.

**B)** Confocal images of hemocytes and fat body from WL *Lsp2Gal4,UAS-GFP;HmldeltaRFP*. The immunolabeling were done with anti-Srp (in gray), anti-RFP (in red), anti-GFP (in green) and the nuclei were marked with DAPI (blue). Full stacks are displayed, the left panels show the overlay of all channels and then from left to right, single channel are displayed (RFP, Srp and GFP). The scale bars represent 50 μm.

**C)** FACS analysis of the levels of GFP in the circulating and resident hemocytes from *Lsp2Gal4,UAS-GFP;HmldeltaRFP* WL. The three first panels indicate how single cells were selected and the last panels represent the distribution of the cells according to the GFP levels. Note the strong enrichment of GFP positive cells in the circulating pool (in green) as compared to the resident pool (in red).

**D)** Dot plot of genes enriched in the PL-ImpL2 cluster in the NI dataset.

**E)** Confocal images of hemocytes from *GstD-LacZ* wandering larvae. The immunolabeling were done with anti-Srp (in green), anti-LacZ (in gray), phalloidin-Cy3 (in red) and the nuclei were marked with DAPI (blue). Full stacks are displayed, the top panel show the overlay of all channels, the middle panel show the overlay Srp/phalloidin and the lower panel LacZ alone. The scale bar represents 10 μm. Note that this field displaying two LacZ positive cells was selected for containing these cells but is not representative of the whole population of hemocytes, which is largely LacZ negative.

**F)** Phagocytosis assay on WL hemocytes with FITC-beads (in green). Hemocytes were labelled with anti-NimC1 (in gray) and anti-Srp (in red) and the nuclei were marked with DAPI. The images are full stack from confocal acquisition, the scale bar represents 20 μm. The images were acquired after 2 min of incubation with the beads. The percentage of cells containing beads as a function of the time of incubation is represented on the right panel.

**Figure S6: Description of the wasp infestation conditions**

**A)** Time-line of larval development in normal condition and upon wasp infestation. The infestation conditions and the collection time point are indicated.

**B,C)** Status of the lymph gland (intact or histolysed) from mid L2 stage to wandering larval stage in normal condition and upon wasp infestation. The primary and secondary lobes are indicated by white dashed lines. The scale bars represent 100 μm **(B)**. The rate of histolysis during larval development was quantified in **(C)**. The x-axis represents the time-line in h AEL and the y-axis the proportion of lymph glands displaying histolysis. Wasp infestation is done at 54 h AEL.

**D)** Number of PL (in red) and LM (in green) after wasp infestation. Hemocytes were counted at 54 h AEL (time of infestation, L2), 78 h AEL (early L3), 102 h AEL (mid L3) and 116 h AEL (WL). The number of hemocytes counted in control WL is indicated with a gray dash line.

**E)** Correlation (Pearson) between the clusters from the NI dataset (vertical) and the WI dataset (horizontal). The Pearson score is represented with a color gradient from blue to white to red for strong correlation. Note the strong correlation between NI and WI for each cluster (red diagonal).

**Figure S7: origin of the hemocyte populations in NI and WI wandering larvae.**

**A,B)** Immunolabeling of lymph glands from WL *UAS-FLP/+;srp(hemo)Gal4/gtrace-mCD8-GFP;gtrace-nls-GFP/+* **(A)** and *UAS-FLP/+;DotGal4/gtrace-mCD8-GFP;gtrace-nls-GFP/+* **(B)** with anti-GFP (in green) and the nuclei are labelled with DAPI. The scale bars represent 100 μm. The primary and secondary lobes are highlighted by white dashed lines. Note that with the driver *srp(hemo)Gal4* **(A)**, the GFP signal is completely absent from the lymph gland lobes, while with the *DotGal4* **(B)**, all the cells from the lobes are GFP +.

**C,D)** Immunolabeling of hemocytes from WL *UAS-FLP/+;srp(hemo)Gal4/gtrace-mCD8-GFP;gtrace-nls-GFP/+* **(C)** and *UAS-FLP/+;DotGal4/gtrace-mCD8-GFP;gtrace-nls-GFP/+* **(D)** with anti-GFP (in green) and the nuclei are labelled with DAPI. The scale bars represent 10 μm. Note that with the driver *srp(hemo)Gal4* **(C)**, most hemocytes are labelled with GFP, while with the *DotGal4* **(D)**, GFP is absent from the hemocytes.

**E,F)** Immunolabeling of hemocytes from WL *UAS-FLP/+;srp(hemo)Gal4/gtrace-mCD8-GFP;gtrace-nls-GFP/+* **(E)** and *UAS-FLP/+;DotGal4/gtrace-mCD8-GFP;gtrace-nls-GFP/+* **(F) after wasp infestation** with anti-GFP (in green) and the nuclei are labelled with DAPI. The scale bars represent 10 μm. Note that GFP PL and LM are observed with the two drivers, indicating that the hemocytes from the two hematopoietic waves coexist in circulation in the hemolymph of WI larvae.

**Figure S8: Metabolic pathways in the single cell RNA seq NI and WI clusters.**

Dot plots of genes of the different metabolic pathways in the clusters from the single cell datasets (NI in blue, WI in red).

### Supplemental Tables

**Table S1: comparison of the bulk RNA seq from E16 and WL and GO term enrichment analysis**

The file contains two tables. The first one (Bulk E16 vs WL) contains the results of the comparison of the transcriptomes from E16 and WL hemocytes by DeSEQ2. The table indicates the gene id (col 1), the base mean determined by DeSEQ2 (col 2), the expression levels in E16 and WL (col 3 and 4), the fold change and the log2 fold change WL/E16 (col 5 and 6), the p-value and the adjusted p-value (col 7 and 8) and the annotation as extracellular matrix gene (col 9). The second table (GO term enrichment analysis) contains the results of the GO term enrichment analysis carried out on the E16 and WL hemocyte transcriptomes. The analysis was done on the genes enriched in WL, enriched in E16 and common between the two stages (indicated in col 1). The table indicates also the category of the GO term (col 2), the GO term (col 3), the number of genes associated with the GO term (col 4), the p-value of the GO term enrichment (col 5) and the enrichment of the GO term (col 6).

**Table S2: list of the significant markers for each cluster of the NI dataset**

The table contains the gene symbol (col 1), the gene Fbgn ID (col 2), the average log fold change compared to the other clusters (col 3), the percentage of cells of the cluster expressing the marker (col 4), the percentage of cells of the whole dataset expressing the marker (col 5), the p-value and the adjusted p-value (col 6 and 7) and the cluster in which the marker is expressed (col 8).

**Table S3: GO terms enriched in the clusters from the NI dataset**

The table indicates the NI cluster (col 1), the category of the GO term (col 2), the GO term (col 3), the number of genes associated with the GO term (col 4), the p-value of the GO term enrichment (col 5) and the enrichment of the GO term (col 6).

**Table S4: list of genes involved in the metabolic pathways.**

This table is related to the data presented in **Figure S2 and S8.**

For each metabolic pathway, the table indicate the level at which the gene acts (col 1), the name of the enzyme (col 2), the enzyme E.C. number (col 3), the gene symbol (col 4), the Fbgn ID (col 5), the log2 fold change (WL/E16) (col 6), the adjusted p-value (col 8) and the levels and log2 fold change in whole WL and whole E16 from modEncode (col 8, 9 and 10). Enzymes highlighted in orange are significantly up-regulated or down-regulated in the WL and presented in **Figure S2**.

**Table S5: list of the primers used in this study**

The table indicate the gene symbol, the forward primer and the reverse primer.

**Table S6: list of the p-values of Figure 3A, 7A,E**

**Table S7: list of the most significant markers for the clusters (Ni and WI combined) and their orthologues.**

The table indicates the cluster (col 1), the gene symbol (col 2), the gene Fbgn ID (col 3), the AUC score determined with roc test from Seurat (col 4), the average difference compared to all other clusters (col 5) percentage of cells of the cluster expressing the marker (col 4), the percentage of cells of the whole dataset expressing the marker (col 5), the percentage of cell of the cluster expressing the marker (col 6) and the human orthologue of the marker determined with DIOPT (Hu et al., 2011) (col 7, none if no orthologues were found).

